# Development of novel tools for dissection of central versus peripheral dopamine D_2_-like receptor signaling in dysglycemia

**DOI:** 10.1101/2024.02.21.581451

**Authors:** Alessandro Bonifazi, Michael Ellenberger, Zachary J. Farino, Despoina Aslanoglou, Rana Rais, Sandra Pereira, José O. Mantilla-Rivas, Comfort A. Boateng, Amy J. Eshleman, Aaron Janowsky, Margaret K. Hahn, Gary J. Schwartz, Barbara S. Slusher, Amy Hauck Newman, Zachary Freyberg

## Abstract

Dopamine (DA) D_2_-like receptors in both the central nervous system (CNS) and the periphery are key modulators of metabolism. Moreover, disruption of D_2_-like receptor signaling is implicated in dysglycemia. Yet, the respective metabolic contributions of CNS versus peripheral D_2_-like receptors including D_2_ (D2R) and D_3_ (D3R) receptors remain poorly understood. To address this, we developed new pharmacological tools, D_2_-like receptor agonists with diminished and delayed blood-brain barrier capability, to selectively manipulate D2R/D3R signaling in the periphery. We designated bromocriptine methiodide (BrMeI), a quaternary methiodide analogue of D2/3R agonist and diabetes drug bromocriptine, as our lead compound based on preservation of D2R/D3R binding and functional efficacy. We then used BrMeI and unmodified bromocriptine to dissect relative contributions of CNS versus peripheral D2R/D3R signaling in treating dysglycemia. Systemic administration of bromocriptine, with unrestricted access to CNS and peripheral targets, significantly improved both insulin sensitivity and glucose tolerance in obese, dysglycemic mice *in vivo*. In contrast, metabolic improvements were attenuated when access to bromocriptine was restricted either to the CNS through intracerebroventricular administration or delayed access to the CNS via BrMeI. Our findings demonstrate that the coordinated actions of both CNS and peripheral D_2_-like receptors are required for correcting dysglycemia. Ultimately, the development of a first-generation of drugs designed to selectively target the periphery provides a blueprint for dissecting mechanisms of central versus peripheral DA signaling and paves the way for novel strategies to treat dysglycemia.

## Introduction

Dopamine (DA) is increasingly recognized as an important modulator of metabolism^1–13^. Until now, most studies examining dopaminergic modulation of metabolism have primarily focused on DA D_2_-like receptors (D_2_, D_3_ and D_4_ receptors) in brain regions associated with metabolic regulation including striatum and hypothalamus^3, 10, 14–16^. For example, hypothalamic dopamine D_2_ (D2R) and D_3_ (D3R) receptors mediate appetite and feeding behaviors, while blockade of these receptors with antipsychotic drugs (APDs) impairs central glucose sensing^17–22^. Changes in the expression of striatal D_2_-like receptors are also associated with overeating and obesity^23, 24^. Moreover, D2R polymorphisms are associated with insulin resistance and type 2 diabetes (T2D)^25^. Nevertheless, the precise mechanisms and sites of action for DA’s metabolic effects remain unclear.

Discovery of DA D_2_-like receptors outside of the central nervous system (CNS) has expanded the scope of DA’s roles as a metabolic modulator^9, 26–28^. In the endocrine pancreas, D2R and D3R are expressed alongside the DA biosynthetic and catabolic machinery in glucagon-secreting α-cells and insulin-secreting β-cells^9, 11, 13, 27, 29, 30^. We and others demonstrated that α- and β-cells produce DA which then acts locally through islet D2R and D3R to negatively modulate hormone secretion in an autocrine/paracrine manner^9, 11, 13, 27, 29, 31, 32^. Importantly, interfering with peripheral D2R/D3R signaling contributes to dysglycemia. We recently discovered that disruption of pancreatic α-cell and β-cell D2R/D3R signaling by APDs significantly elevates insulin and glucagon secretion – hallmarks of dysglycemia^13^. This is consistent with earlier reports of APD-induced hyperinsulinemia and hyperglucagonemia^33–35^. As in T2D, the increases in circulating insulin and glucagon due to disrupted D2R/D3R signaling may desensitize insulin-sensitive peripheral targets (*e.g.*, liver, skeletal muscle, adipose tissue) and stimulate glucose production over time to ultimately produce sustained hyperglycemia and insulin resistance^26, 36, 37^. Conversely, D2R/D3R stimulation with agonists such as bromocriptine or cabergoline acts on targets in the periphery including pancreatic islets and adipose tissue to improve insulin sensitivity^12, 32, 38–40^. While these drugs can act on other targets (*e.g.*, serotonin and adrenergic receptors), importantly, D2R/D3R agonism appears to be the principal mechanism by which they ameliorate dysglycemia^41^. Indeed, bromocriptine is FDA-approved as a novel therapeutic agent for T2D^42–44^. Taken together, these reports suggest an important role for peripheral D2R/D3R signaling in metabolic regulation and dysglycemia, particularly within the pancreas – observations that have been reproducible across cell, animal, and human studies^26, 28, 37^.

To date, it has been difficult to disentangle the respective metabolic contributions of CNS versus peripheral D_2_-like receptor signaling to dysglycemia and its effective treatment. Initial work to address the metabolic roles of D_2_-like receptors employed transgenic global D2R or D3R knockout (KO) mouse models^10, 15, 45^. However, these D2R or D3R KO mice exhibited complex metabolic phenotypes due to concurrent effects of receptor deletion on both CNS and peripheral targets^15, 45–47^. More recently, mice with constitutive D2R KO specifically in β-cells demonstrated inappropriately elevated serum insulin levels in response to meals, suggesting a key role for β-cell D2R signaling in insulin release *in vivo*^11^. Nevertheless, genetic constitutive KO strategies may still lead to potential compensatory effects at systemic, cellular, and/or transcriptional levels. Such alterations may therefore confound interpretation of metabolic phenotypes, particularly for discerning the relative metabolic contributions of CNS versus peripheral DA signaling.

In lieu of inducible KO models, pharmacological strategies using drugs that selectively stimulate or block DA receptor actions at either CNS or peripheral targets offer an effective alternative approach to dissect central versus peripheral metabolic contributions of DA signaling. Such drug approaches have key advantages since: 1) pharmacological manipulations are reversible, and 2) offer the ability to finely tune changes to receptor signaling either acutely or chronically by controlling drug doses and durations of exposure. Here, we describe the development of the first generation of new pharmacological tools intended to selectively target peripheral D_2_-like receptors. Our aim was to employ peripherally-limited drugs to specifically examine the metabolic relevance of peripheral D_2_-like receptor signaling in dysglycemia and its treatment without the confounds of CNS actions. We used quaternary methiodide (MeI) conjugation as a strategy to diminish blood brain barrier (BBB) permeability, selecting bromocriptine methiodide (BrMeI) as our lead compound. We then tested BrMeI *in vivo*, comparing its actions to those of unmodified bromocriptine, to pharmacologically dissect the relative metabolic contributions of CNS versus peripheral D_2_-like receptors in dysglycemic mice. Systemic administration of bromocriptine, which has access to both CNS and peripheral targets, significantly improved insulin sensitivity and glucose tolerance. In contrast, selectively limiting D2R/D3R agonism either to the CNS (via intracerebroventricular bromocriptine administration) or to the periphery (via BrMeI) abolished metabolic improvements. Together, these results suggest the importance of coordinated, tandem signaling by both peripheral and CNS D_2_-like receptors for glycemic control and offers a new approach both for studying metabolism as well as for the treatment of dysglycemia.

## Materials and Methods

### Chemistry

#### General Information

All chemicals and solvents were purchased from the specified chemical suppliers unless otherwise stated and used without further purification. All melting points were determined on an OptiMelt automated melting point system (Stanford Research Systems, Sunnyvale, CA) and were uncorrected. Nuclear magnetic resonance (NMR) spectra including the ^1^H and ^13^C NMR spectra were recorded on a Varian Mercury Plus 400 NMR spectrometer (Agilent Technologies, Santa Clara, CA). Proton chemical shifts are reported as parts per million (δ ppm) relative to tetramethylsilane (0.00 ppm) as an internal standard. Coupling constants were measured in Hz. Chemical shifts for ^13^C NMR spectra are reported as parts per million (δ ppm) relative to deuterated CHCl_3_ or deuterated MeOH (CDCl_3_, 77.5 ppm, CD_3_OD 49.3 ppm). Gas chromatography-mass spectrometry (GC/MS) data were acquired (where obtainable) using an Agilent Technologies 6890N GC system equipped with an HP-5MS column (cross-linked 5% PH ME siloxane, 30 m × 0.25 mm i.d. × 0.25 μm film thickness) and a 5973 mass-selective ion detector in electron-impact mode. Ultrapure grade helium was used as the carrier gas at a flow rate of 1.2 mL/min. The injection port and transfer line temperatures were 250°C and 280°C, respectively, and the oven temperature gradient was as follows: the initial temperature (100°C) was held for 3 min, increased to 295°C at 15°C/min over 13 min, and maintained at 295°C for 10 min. All column chromatography was performed using Merck silica gel (230-400 mesh, 60Å; Merck, Rahway, NJ) or via preparative thin layer chromatography using Analtech silica gel (1000 µm; Analtech, Newark, DE). Microanalyses were performed by Atlantic Microlab, Inc. (Norcross, GA) and agree within ±0.4% of calculated values. All compounds were evaluated to be >95% pure based on elemental analysis (see Supplementary Table 1), NMR, HRMS (high resolution mass spectroscopy within ±5 ppm agreement), and MS/MS (ESI in positive mode) approaches.

### General procedure for the synthesis of methiodide analogs

A solution of the free base form of the respective parent compounds in acetone or ethanol was added with excess of methyl iodide (3 eq). After 24 h stirring at room temperature in the dark, the solid products were filtered and washed with diethyl ether to yield pure quaternary ammonium salts.

### (4a*R*,8a*R*)-5-Methyl-5-propyl-4,4a,5,6,7,8,8a,9-octahydro-2*H*-pyrazolo[3,4-*g*]quinolin-5-ium iodide (AB01-59, (-)-Quinpirole MeI)

Commercially available (-)-quinpirole HCl (Tocris, Bristol, United Kingdom; 50 mg, 0.19 mmol) was washed with saturated NaHCO_3_ aqueous solution and extracted with dichloromethane (DCM). The organic layer was dried with Na_2_SO_4_, filtered, and evaporated to yield the free base. The methiodide derivative was prepared according to the general procedure. ^1^H NMR (400 MHz, CD_3_OD) δ 1.05 (t, *J* = 7.2 Hz, 3H), 1.53 (m, 1H), 1.82-1.92 (m, 3H), 2.05-2.15 (m, 2H), 2.43-2.53 (m, 2H), 2.85 (m, 1H), 3.03 (dd, *J* = 4.4, 4.4 Hz, 1H), 3.12 (s, 3H), 3.39-3.70 (m, 6H), 7.46 (br s, 1H). HRMS ESI-MS [M]^+^ found = 234.1964; calculated = 234.1965. [α]^24^_D_ = −65.83° (CH_3_OH, *c* = 0.12). m.p. 211-213°C. Anal. (C_14_H_24_IN_3_·0.5H_2_O) C, H, N. Presence of di-methylated analog (C_15_H_26_IN_3_) observed in HRMS ESI-MS [M]^+^ found = 248.2118.

(6a*R*,9*R*)-5-Bromo-9-(((2*R*,5*S*,10a*S*,10b*S*)-10b-hydroxy-5-isobutyl-2-isopropyl-3,6-dioxooctahydro-8*H*-oxazolo[3,2-*a*]pyrrolo[2,1-*c*]pyrazin-2-yl)carbamoyl)-7,7-dimethyl-4,6,6a,7,8,9-hexahydroindolo[4,3-*fg*]quinolin-7-ium iodide (AB01-60, Bromocriptine MeI) was prepared according to the general procedure starting from commercially available bromocriptine free base (Santa Cruz Biotechnology, Dallas, TX; 150 mg, 0.23 mmol).^1^H NMR (400 MHz, CD_3_OD) δ 0.75 (d, *J* = 6.4 Hz, 3H), 1.00 (dd, *J* = 6.4, 6.4 Hz, 6H), 1.18 (d, *J* = 6.8 Hz, 3H), 1.75-2.11 (m, 7H), 2.27 (m, 1H), 3.12 (dd, *J* = 12.4, 12 Hz, 1H), 3.30 (br s, 3H), 3.46-3.53 (s and m, 3H and 2H), 3.61 (dd, *J* = 5.6, 6.0 Hz, 1H), 3.80 (dd, *J* = 6.0, 6.0 Hz, 1H), 3.91-4.04 (m, 2H), 4.15-4.19 (m, 1H), 4.49 (dd, *J* = 5.2, 5.6 Hz, 1H), 4.71-4.76 (m, 1H), 6.67 (br s, 1H), 7.16 (t, *J* = 7.8 Hz, 1H), 7.24 (d, *J* = 7.6 Hz, 1H), 7.29 (d, *J* = 7.2 Hz, 1H). ^13^C NMR (100 MHz, CD_3_OD) δ 14.69, 15.83, 20.52, 20.85, 21.51, 21.87, 24.60, 25.80, 33.46, 33.50, 43.33, 45.93, 53.20, 62.64, 64.15, 68.68, 91.17, 104.00, 150.03, 150.60, 110.41, 113.21, 116.29, 122.90, 123.11, 125.20, 130.31, 134.68, 166.04, 166.34, 171.87. HRMS ESI-MS [M]^+^ found = 668.2449; calculated = 668.2442. [α]^24^_D_ = +75.00° (CH_3_OH, *c* = 0.32). m.p. 209-210°C. Anal. (C_33_H_43_BrIN_5_O_5_) C, H, N.

**4-(2,3-Dichlorophenyl)-1-methyl-1-(4-((2-oxo-1,2,3,4-tetrahydroquinolin-7-yl)oxy)butyl)piperazin-1-ium iodide (AB01-61, Aripiprazole MeI)** was prepared according to the general procedure, starting from commercially available aripiprazole free base (Sigma Aldrich, Saint Louis, MO; 50 mg, 0.11 mmol).^1^H NMR (400 MHz, CD_3_OD) δ 1.90-1.93 (m, 2H), 2.05-2.09 (m, 2H), 2.53 (t, *J* = 7.6 Hz, 2H), 2.86 (t, *J* = 7.6 Hz, 2H), 3.26 (s, 3H), 3.44 (m, 4H), 3.62-3.71 (m, 6H), 4.07 (t, *J* = 5.8 Hz, 2H), 6.49 (d, *J* = 2.4 Hz, 1H), 6.59 (dd, *J* = 2.8, 2.4 Hz, 1H), 7.07 (d, *J* = 8.0 Hz, 1H), 7.23-7.32 (m, 3H). m.p. 133°C (decomposition). Anal. (C_24_H_30_Cl_2_IN_3_O_2_) C, H, N.

#### 7-Hydroxy-N-methyl-N,N-dipropyl-1,2,3,4-tetrahydronaphthalen-2-aminium iodide (AB01-62, 7-OH-DPAT MeI)

Commercially available (±)-7-hydroxy-DPAT HBr (Tocris; 50 mg, 0.15 mmol) was washed with saturated NaHCO_3_ aqueous solution and extracted with DCM. The organic layer was dried with Na_2_SO_4_, filtered, and evaporated to yield the free base. The methiodide derivative was prepared according to the general procedure. ^1^H NMR (400 MHz, CD_3_OD) δ 1.04 (t, *J* = 7.0 Hz, 6H), 1.76-1.83 (m, 4H), 1.90-1.95 (m, 1H), 2.40 (m, 1H), 2.82-3.07 (m, 2H), 3.11 (s, 3H), 3.20 (m, 1H), 3.24-3.48 (m, 4H), 3.84 (m, 1H), 6.60 (m, 2H), 6.95 (d, *J* = 8.4 Hz, 1H). m.p. 126-127°C. Anal. (C_17_H_28_INO·0.25H_2_O) C, H, N.

### (*R*)-*N*,*N*,*N*-Trimethyl-2-oxo-1,2,5,6-tetrahydro-4*H*-imidazo[4,5,1-*ij*]quinolin-5-aminium iodide (MPE01-05, Sumanirole MeI)

Sumanirole maleate (230 mg, 1.1 mmol) was synthesized as described previously^48, 49^. The compound was washed with saturated NaHCO_3_ aqueous solution and extracted with DCM. The organic layer was dried with Na_2_SO_4_, filtered, and evaporated to yield the free base. The methiodide derivative was prepared according to the general procedure.^1^H NMR (400 MHz, CD_3_OD) δ 3.23 (br s, 9H), 3.36 (m, 1H), 3.82-4.20 (m, 3H), 4.42 (dd, *J* = 2.8, 2.4 Hz, 1H), 6.85-7.15 (m, 3H).^13^C NMR (100 MHz, CD_3_OD) δ 23.86, 24.60, 27.72, 37.62, 40.77, 40.47, 51.93, 68.04, 107.68, 107.80, 113.74, 114.44, 119.51, 119.87, 122.12, 126.25, 126.81, 154.68. ESI-MS [M]^+^ = 232.11. [α]^24^_D_ = −8.33° (CH_3_OH, *c* = 0.24). m.p. 180°C (decomposition). Anal. (C_13_H_18_IN_3_O·H_2_O) C, H, N.

**1-((1*H*-Indol-3-yl)methyl)-4-(4-chlorophenyl)-4-hydroxy-1-methylpiperidin-1-ium iodide (MPE01-06, L741,626 MeI)** was prepared according to the general procedure, starting from L741,626 free base, prepared as previously described^50^ (500 mg, 1.5 mmol).^1^H NMR (400 MHz, CD_3_OD) δ 1.87 (br d, *J* = 14.8 Hz, 2H), 2.42 (td, *J* = 4.0, 2.4, 4.4 Hz, 2H), 3.20 (s, 3H), 3.26-3.31 (m, 2H), 3.43 (dd, *J* = 2.0, 2.0 Hz, 2H), 3.87 (td, *J* = 2.8, 2.8, 2.8 Hz, 2H), 7.17-7.24 (m, 2H), 7.35 (dd, *J* = 2.0, 2.0 Hz, 2H), 7.47-7.59 (m, 3H), 7.70 (s, 1H), 7.80 (d, *J* = 5.6 Hz, 1H). ^13^C NMR (100 MHz, CD_3_OD) δ 32.29, 43.27, 55.15, 63.77, 67.53, 100.71, 110.96, 117.79, 118.04, 118.26, 118.79, 120.48, 121.28, 122.17, 124.48, 126.10, 126.32, 127.82, 128.06, 128.13, 130.09, 132.88, 134.61, 145.21. m.p. 177-178°C. Anal. (C_21_H_24_ClIN_2_O) C, H, N.

### (4*S*,4a*R*,10b*R*)-9-Hydroxy-4-methyl-4-propyl-3,4,4a,10b-tetrahydro-2*H*,5*H*-chromeno[4,3-*b*][1,4]oxazin-4-ium iodide (AB01-117, (+)-PD128,607 MeI)

Commercially available (+)-PD128,607 HCl (Tocris or OxChem, Wood Dale, IL; 100 mg, 0.15 mmol) was washed with 28% NH_4_OH aqueous solution and extracted with DCM. The organic layer was dried with Na_2_SO_4_, filtered, and evaporated to yield the free base. The methiodide derivative was prepared according to the general procedure.^1^H NMR (400 MHz, d_6_-DMSO) δ 0.91 (t, *J* = 7.2 Hz, 3H), 1.74 (m, 2H), 3.13 (s, 3H), 3.29-3.72 (m, 4H), 3.84 (t, *J* = 9.8 Hz, 1H), 4.02-4.26 (m, 3H), 4.76 (d, *J* = 8.4 Hz, 1H), 5.28 (d, *J* = 9.6 Hz, 1H), 6.61-6.73 (m, 3H), 9.13 (br s, 1H). ^13^C NMR (100 MHz, CD_3_OD) δ 10.92, 15.00, 31.16, 41.18, 59.88, 61.00, 65.83, 66.74, 68.64, 100.55, 100.56 112.30, 117.07, 117.12 121.36, 145.40, 152.14. HRMS ESI-MS [M]^+^ found = 264.15911; calculated = 264.15942. [α]^23^_D_ = +56.8° (CH_3_OH, *c* = 0.19). m.p.= 160-162°C. Anal. (C_15_H_22_INO_3_.1/3NH_4_Cl.1/3H_2_O) C, H, N.

**1-Methyl-1-(2-oxo-2-(*m*-tolylamino)ethyl)-4-phenylpiperidin-1-ium iodide (AB01-102, CAB03-015 MeI)** was prepared according to the general procedure, starting from CAB03-015 (100 mg, 0.3 mmol) as previously described^51^. ^1^H NMR (400 MHz, CD_3_CN) (mixture of stereoisomers) δ 2.04-2.30 (m, 4H), 2.35 (s, 3H), 2.95 (m, 1H), 3.40 (s, 3H), 3.43 (m, 1H), 3.74-3.88 (m, 2H), 4.03 (br d, *J* = 12.4 Hz, 1H), 4.47 (br s, 2H, isomer 75%), 4.59 (br s, 2H, isomer 25%), 7.03 (d, *J* = 6.4 Hz, 1H), 7.28-7.58 (m, 8H), 9.66 (br s, 1H, isomer 75%), 9.83 (br s, 1H, isomer 25%). ^13^C NMR (100 MHz, CD_3_CN) δ 20.55, 26.81, 27.18, 38.68, 39.06, 45.72, 54.02, 57.38, 62.03, 62.82, 65.27, 67.09, 99.90, 120.58, 122.06, 125.70, 125.76, 126.75, 126.96, 127.27, 128.59, 128.69, 128.77, 128.80, 133.36, 137.58, 138.92, 143.17, 143.27, 161.11. m.p. 57-58°C. Anal. (C_21_H_27_IN_2_O) C, H, N.

### Radioligand binding studies

#### Cell culture and sample preparation

HEK293 cells stably expressing human D2R, D3R, or D4R were grown in a 50:50 mix of DMEM and Ham’s F12 culture media supplemented with 20 mM HEPES, 2 mM L-glutamine, 0.1 mM non-essential amino acids, 1x antibiotic/antimycotic, 10% heat-inactivated fetal bovine serum (FBS), and 200 µg/ml hygromycin (Life Technologies, Grand Island, NY) (37°C, 5% CO_2_). Upon reaching 80-90% confluence, cells were harvested using pre-mixed Earle’s Balanced Salt Solution (EBSS) with 5 mM EDTA (Life Technologies) and centrifuged at 3000 rpm (10 min, 21°C). Following removal of the supernatant, cell pellets were resuspended in 10 mL hypotonic lysis buffer (5 mM MgCl_2_, 5 mM Tris, pH 7.4 at 4°C) and centrifuged at 20000 rpm (30 min, 4°C). Pellets were subsequently resuspended in fresh binding buffer, with protein concentrations determined via Bradford protein assays (Bio-Rad, Hercules, CA).

#### Radioligand competition binding

The radioligand [^3^H]-(R)-(+)-7-OH-DPAT was employed in all competition binding studies with the exceptions of L741,626 and its MeI analogue, MPE01-06, which instead used [^3^H]-*N*-methylspiperone. For [^3^H]-(R)-(+)-7-OH-DPAT binding studies, membranes were harvested fresh; the binding buffer was made from 50 mM Tris, 10 mM MgCl_2_, 1 mM EDTA, pH 7.4. On test day, all test compounds were freshly dissolved in 30% DMSO and 70% H_2_O to a stock concentration of 1 mM or 100 µM. For [^3^H]-*N*-methylspiperone binding studies, the binding buffer was composed of 8.7 g/L Earle’s Balanced Salts without phenol red (US Biological, Salem, MA), 2.2 g/L sodium bicarbonate, pH 7.4; 500 µg/ml membranes were stored at −80°C until use. To assist in the solubilization of free-base compounds, 10 µL of glacial acetic acid was added along with DMSO. Each test compound was serially diluted using 30% DMSO vehicle. Membranes were diluted in fresh binding buffer. Radioligand competition experiments were conducted in 96-well plates containing 300 µL fresh binding buffer, 50 µL of diluted test compound, 100 µL of membranes ([^3^H]-*N*-methylspiperone: 20 µg total protein for hD2R or hD3R; [^3^H]-(R)-(+)-7-OH-DPAT: 80 µg, 40 µg and 60 µg total protein for hD2R, hD3R or hD4R respectively), and 50 µl of radioligand diluted in binding buffer ([^3^H]-*N*-methylspiperone: 0.4 nM final concentration, Perkin Elmer, Waltham, MA; [^3^H]-(R)-(+)-7-OH-DPAT: 1.5 nM final concentration for hD2R, 0.5 nM final concentration for hD3R and 3 nM final concentration for hD4R; ARC, Saint Louis, MO). Nonspecific binding was determined using 10 µM (+)-butaclamol (Sigma-Aldrich) and total binding was determined with 30% DMSO vehicle. All compound dilutions were tested in duplicate or triplicate with reactions incubated at room temperature for either 60 min in [^3^H]-*N*-methylspiperone studies or for 90 min in [^3^H]-(R)-(+)-7-OH-DPAT studies. Reactions were terminated by filtration through a Perkin Elmer Uni-Filter-96 GF/B 96-well microplate presoaked in 0.5% polyethylenimine using a Brandel 96-Well Plate Harvester Manifold (Brandel Instruments, Gaithersburg, MD). The filters were washed 3 × 1 mL/well of ice-cold binding buffer. Perkin Elmer MicroScint 20 Scintillation Cocktail (65 µL/well) was added and filters were counted using a Perkin Elmer MicroBeta Microplate Counter. IC_50_ values for each compound were determined from dose-response curves, enabling calculation of K_i_ values using the Cheng-Prusoff equation^52^. Analyses of receptor binding data were performed using GraphPad Prism (version 6.0; GraphPad Software, San Diego, CA). K_i_ values for each compound were determined from at least three independent experiments performed in triplicate. Receptor binding data for (-)-quinpirole and sumanirole parent compounds were previously reported^49^.

### Mitogenesis assays

DA D_2_-like receptor-mediated mitogenesis functional assays were conducted as reported earlier^48,53^. Briefly, Chinese hamster ovary (CHO) cells expressing human D2R or D3R (CHOp-D2R or CHOp-D3R, respectively) were maintained in α-MEM culture media supplemented with 10% FBS, 0.05% penicillin/streptomycin, and 400 µg/mL geneticin (G418) (37°C, 5% CO_2_). To measure D_2_-like receptor-mediated stimulation of mitogenesis via D2R or D3R, the respective cell lines were seeded into 96-well plates at a concentration of 5000 cells/well. After 48-72 h, cells were rinsed twice with serum-free α-MEM and incubated for an additional 24 h (37°C, 5% CO_2_). On test day, serial dilutions of the test compounds were made in serum-free α-MEM media. Medium was then replaced with 100 μL of the respective test compound dilutions. Following ∼24 h of drug incubation (37°C, 5% CO_2_), 0.25 μCi of [^3^H]thymidine in α-MEM supplemented with 10% FBS was added to each well and incubated for an additional 2 h (37°C, 5% CO_2_). At the assay conclusion, cells were trypsinized (1% trypsin in Ca^2+^/Mg^2+^-free PBS). Plates were then filtered, and intracellular radioactivity was measured via scintillation spectrometry. The D2R/D3R agonist (-)-quinipirole was run each day as an internal control. Specific [^3^H]thymidine incorporation at each drug dose was expressed as % maximal quinpirole effect for each of the tested drugs. EC_50_ values and *E_max_* values for each compound were determined from dose-response curves via nonlinear regression analyses using GraphPad Prism software.

### Dopamine D_4_ receptor-mediated adenylate cyclase activity/cAMP functional assay

HEK-D4.4-AC1 cells expressing human DA D4.4 receptors and adenylate cyclase type I were used. Experiments were conducted with a cAMP enzyme immunoassay (EIA) kit (Cayman, Ann Arbor, MI, USA) as described previously^54^. Basal cAMP was subtracted from all cAMP values. Maximal DA D4.4 receptor-mediated inhibition of forskolin-stimulated cAMP formation by the agonist drugs was defined with 1 µM (-)-quinpirole. The maximal effect of the assayed drugs was then normalized to maximal effect of (-)-quinpirole.

#### Glucose-stimulated insulin secretion assay Cell culture

INS-1E cells (gift of Dr. Pierre Maechler, Université de Geneve) were cultured as described earlier^11, 27, 55^. Cells were maintained in RPMI 1640 medium supplemented with 5% heat-inactivated FBS, 2 mM L-glutamine, 10 mM HEPES, 1 mM sodium pyruvate, 100 U/mL penicillin/streptomycin, and 50 µM 2-mercaptoethanol (37°C, 5% CO_2_).

#### Glucose-stimulated insulin secretion (GSIS)

Assays measuring GSIS in INS-1E cells were conducted as reported earlier^11, 56^. Briefly, INS-1E cells were seeded into a 24-well plate at an initial seeding density of 5.0×10^5^ cells/well. RPMI 1640 media was exchanged 24 h following plating and experiments were conducted the next day. On the experimental day, cells were glucose-starved (1 h, 37°C) in KRB and subsequently stimulated with 20 mM glucose in the presence of drugs (90 min, 37°C). At assay conclusion, supernatants were collected for insulin detection.

#### Insulin measurement and analysis

Secreted insulin was detected using a commercially available insulin detection kit (PerkinElmer/Cisbio Bioassays, Bedford, MA) based on homogenous time-resolved fluorescence resonance energy transfer (HTRF) technology as described previously^11, 56, 57^. Samples were incubated with the antibodies for 2 h at room temperature and then read using a PheraStar FS multi-mode plate reader (BMG Labtech, Ortenberg, Germany). Insulin concentrations (in ng/mL) were derived via extrapolation of ratiometric fluorescence readings (665 nm/620 nm) to a second-order quadratic polynomial curve. Dose-response curves were fit via non-linear regression of Log[drug] versus normalized % maximum insulin secretion via GraphPad software (version 6.0). IC_50_ and *E_max_* values were calculated from the non-linear regression analyses.

### NanoBRET

#### DNA constructs

NanoBRET experiments employed human D2R (DRD2) cDNA tagged at the N-terminus with an IL6 signal sequence followed by a HiBiT tag and tagged at the C-terminus with a HaloTag: IL6-HiBiT-D2R-HaloTag. G protein recruitment studies employed human Gα_i1_ with nanoluciferase (NanoLuc) inserted at position 91: NanoLuc-Gα_i1(91)_. β-arrestin2 recruitment assays used human β-arrestin2 (ARRB2) fused to NanoLuc at its N-terminus: NanoLuc-β-arrestin2. All constructs were prepared by Genscript USA (Piscataway, NJ) and cloned into a pcDNA3.1(+) vector backbone (Thermo Fisher Scientific, Waltham, MA). Constructs were confirmed by sequencing analysis.

#### Transfection

HEK-293T cells were transfected after reaching 70% confluency via Lipofectamine 3000 (Thermo Fisher Scientific) (2.5 μg total cDNA) per manufacturer instructions. We used a 100 (IL6-HiBiT-D2R-HaloTag):1 (NanoLuc-Gα_i1(91)_ or NanoLuc-β-arrestin2) nanoBRET donor/acceptor pair ratio. Empty pcDNA3.1(+) vector was used to maintain a constant amount of total transfected DNA.

#### NanoBRET

NanoBRET assays were conducted as described previously^12^. After transfection, cells were plated onto white poly-D-Lysine pre-coated, 96-well, flat bottom plates (Greiner Bio-One, Monroe, NC) at a density of 5 × 10^4^ cells/well. After adhering to the plates overnight, cells were washed with HBSS and labeled with 100 nM HaloTag NanoBRET 618 ligand (Promega Corp., Fitchburg, WI) in phenol red-free Opti-MEM I reduced serum medium (Gibco/Thermo Fisher Scientific) (2 h, 37°C). Cells were washed with HBSS and 5 mM furimazine was added to each well followed by treatment with the respective drugs. Plates were read within 5-10 min after drug addition on a PHERAstar FSX plate reader (BMG Labtech). The nanoBRET ratio was calculated as the emission of the acceptor (618 nm) divided by the emission of the donor (460 nm). Net nanoBRET values were obtained by subtracting the background nanoBRET ratio from cells expressing only NanoLuc. Data were normalized to the % maximum DA response. NanoBRET data were further normalized to define the minimum and maximum response to the corresponding endogenous ligand. EC_50_ values were calculated by non-linear regression analysis via GraphPad software.

### Initial Screening: Receptor and biogenic amine transporter binding assays

#### General

Test drug BrMeI was weighed and dissolved in DMSO to make a 10 mM stock solution. An initial dilution to 50 µM for binding was made with water (for biogenic amine transporter and DA receptor binding assays) or buffer (for serotonin and opioid receptor binding assays). Subsequent dilutions were made with assay buffer, maintaining a final 0.1% DMSO concentration. Assays for specific binding to D1R, D4R, 5-HT_1A_, 5-HT_2A_, and 5-HT_2C_ were performed as previously described^53^.

#### Data analysis

Excel software (Microsoft Inc., Redmond, WA) was used to analyze data. Binding data were normalized to % inhibition of specific control binding. Nonspecific binding was subtracted from all data. Specific binding in the presence of drug was normalized to binding in the absence of drug (control binding).

### Dopamine receptor binding assays

#### [^3^H]SCH-23390 D1R binding assay

Mouse fibroblast cells expressing human D1R at high density (LhD1 cells) were used. Cells were grown to confluence in DMEM containing 10% FetalClone1 serum (FCS; HyClone, Logan, UT), 0.05% penicillin/streptomycin, and 400 µg/mL G418. One confluent 150 mm plate yielded sufficient membranes for 96 wells with ∼10-15 µg protein/well. Cells were collected via scraping and centrifuged at 500 x *g* for 5 min. The pellet was overlaid with 2 mL assay buffer (50 mM Tris-HCl, 120 mM NaCl, 5 mM KCl, 2 mM CaCl_2_, 1 mM MgCl_2_) and stored at −70°C until use. On the experimental day, the pellet was homogenized in assay buffer with a polytron. Cell homogenate (100 µL) was added to wells containing 800 µL of BrMeI or buffer. After 10 min preincubation, 100 µL of [^3^H]SCH-23390 radioligand (0.18 nM final concentration) was added and incubated at 25°C for 60 min. Reactions were terminated by filtration using a Tomtec 96-well harvester (Tomtec, Hamden, CT) and filter radioactivity was counted using a Perkin Elmer MicroBeta scintillation counter. Nonspecific binding was determined with 1 µM SCH-23390.

#### [^3^H]YM-09151-2 D2R and D3R binding assays

CHO cells expressing human D2R or D3R receptor (CHOp-D2 or CHOp-D3 cells, respectively) were used. Cells were grown to confluence in α-MEM culture medium containing 10% FBS (Atlas Biologicals, Fort Collins, CO), 0.05% penicillin/streptomycin, and 400 µg/mL G418. Membranes were prepared according to the procedures described for D1R-expressing cells using D2R/D3R binding buffer (50 mM Tris containing 120 mM NaCl, 5 mM KCl, 1.5 mM CaCl_2_, 4 mM MgCl_2_, 1 mM EDTA, pH 7.4). One confluent 150 mm plate of D2R-expressing cells, resuspended in 10 mL of binding buffer, yielded enough membranes for 96 wells with ∼10-15 µg protein/well; a single plate of D3R-expressing cells, resuspended in 15 mL, yielded sufficient membranes for 144 wells with ∼7-10 μg protein/well. Cell homogenate (100 µL) was added to wells containing 800 µL of BrMeI or buffer. After 10 min, 100 µL of [^3^H]YM-09151-2 radioligand (0.2-0.5 nM final concentration) was added and incubated at 25°C for 60 min. Reactions were stopped by filtration through polyethylenimine-soaked (0.05%) filters using a Tomtec 96-well harvester. Radioactivity on the filters was counted using a Perkin Elmer MicroBeta scintillation counter. Nonspecific binding was determined with 1 µM chlorpromazine.

#### [^3^H]YM-09151-2 D4R binding assay

Human embryonic kidney (HEK) cells co-expressing the human D4.4 receptor and adenylate cyclase type I (HEK-D4.4-AC1) were used. Cells were grown in DMEM supplemented with 5% FCS, 5% bovine calf serum (BCS), 0.05% penicillin/streptomycin, 2 µg/mL puromycin, and 200 µg/mL hygromycin. One confluent 150 mm plate yielded membranes sufficient for 144 wells with ∼7-10 µg protein/well. Cells were scraped and centrifuged at 500 x *g* for 5 min. The pellet was overlaid with 2 mL binding buffer (50 mM Tris, 120 mM NaCl, 5 mM KCl, 1.5 mM CaCl_2_, 4 mM MgCl_2_, 1 mM EDTA, pH 7.4) and stored at −70°C until use. On the experimental day, the pellet was homogenized in assay buffer with a polytron. Cell homogenate (100 µL) was added to wells containing 800 µL of either BrMeI or binding buffer. After 10 min, 100 µL [^3^H]YM-09151-2 (0.2-0.3 nM final concentration) was added. Plates were incubated at 25°C for 60 min. Reactions were terminated by filtration through polyethylenimine-soaked (0.05%) filters via a Tomtec 96 well harvester. Filter radioactivity was counted using a Perkin Elmer MicroBeta scintillation counter. Nonspecific binding was determined with 1 µM haloperidol.

#### DA receptor binding standards

Specificity of radioligand binding at the respective dopamine receptors was measured via non-radiolabeled standards. For D1R binding assays, inhibition of specific [^3^H]SCH-23390 receptor binding was measured using cold SCH-23390 (100 nM, 10 µM). For D2R and D3R binding assays, inhibition of specific [^3^H]YM-09151-2 receptor binding was measured with cold butaclamol (100 nM, 10 µM). For D4R binding assays, inhibition of specific [^3^H]YM-09151-2 receptor binding was measured using cold haloperidol (100 nM, 10 µM). Inhibition of specific DA receptor binding by the respective standards is summarized in Supplementary Table 5.

### Serotonin receptor binding assays

#### [^3^H]8-OH-DPAT 5-HT_1A_ receptor binding assay

HEK cells expressing the human 5-HT_1A_ receptor (HEK-h5HT1a) were used. Cells were grown to confluence on 150 mm plates in DMEM containing 10% FCS (HyClone), 0.05% penicillin/streptomycin, and 300 µg/mL G418. Cells were collected via scraping into phosphate-buffered saline (PBS) and centrifuged at 270 x *g* for 10 min. The cell pellet was subsequently homogenized in 50 mM Tris-HCl (pH 7.7) with a polytron, and centrifuged at 27000 × *g*. The homogenization and centrifugation were repeated to wash any remaining serotonin (5-HT) from the growth media. The final pellet was resuspended at 0.5 mg protein/mL in assay buffer (25 mM TrisHCl, 100 µM ascorbic acid, 10 µM pargyline, pH 7.4). The assay was performed in triplicate in a 96-well plate. The reaction mixture contained BrMeI, 100 µL of cell homogenate (0.05 mg protein/well) and 100 µL of [^3^H]8-OH-DPAT (0.5 nM final concentration, 170 Ci/mmol, Perkin Elmer) in a final volume of 1 mL. Plates were incubated at room temperature for 60 min and then filtered through polyethylenimine-soaked (0.05%) filters on a Tomtec cell harvester. Filters were then washed with cold 50 mM Tris buffer (pH 7.7) for 6 sec, dried, spotted with scintillation cocktail, and counted for 2 min after a 4 hr delay on a Perkin Elmer Betaplate 1205 liquid scintillation counter (Perkin Elmer). Nonspecific binding was determined with 1.0 µM dihydroergotamine.

#### [^125^I]DOI 5-HT_2A_ and 5-HT_2C_ receptor binding assays

This assay was adapted from previously described studies^58^. Briefly, HEK cells expressing the human 5-HT_2A_ receptor (HEK-h5HT2A) or human 5-HT_2C_ receptor (HEK-h5HT2C) were used. The cells were grown until confluency on 150 mm plates. Cells were washed with PBS, scraped into 2 mL PBS, and stored at −20°C until use. On the experimental day, the cell suspension was thawed and 10 mL assay buffer (50 mM Tris, pH 7.4 at 37°C, with 0.1% ascorbic acid and 5 mM CaCl_2_) was added. This was followed by homogenization via polytron for 5 sec. The homogenate was centrifuged at 15500 rpm for 20 min. To minimize the residual 5-HT concentration, the pellet was resuspended in buffer, homogenized by polytron, and centrifuged as above. The final pellet was resuspended in 10 mL buffer/plate. For the radioligand binding studies, the binding assay included 50 µL BrMeI, 5-HT or buffer, 50 µL cell homogenate, 25 µL [^125^I]DOI (∼0.05 nM) and buffer (final volume = 250 µL). Specific binding was defined as the difference between total binding and binding in the presence of 10 µM 5-HT. The reaction was incubated for 1 hr at 37°C and terminated by filtration thru Perkin Elmer A filtermats presoaked in 0.05% polyethylenimine using a Tomtec 96-well harvester. Remaining filter radioactivity was counted via a Perkin Elmer Betaplate 1205 liquid scintillation counter.

#### 5-HT receptor binding standards

Specificity of radioligand binding at the respective 5-HT receptors was measured via non-radiolabeled standards. For 5-HT_1A_ receptor binding assays, inhibition of specific receptor binding was measured using cold WAY 100,635 (100 nM, 10 µM). For 5-HT_2A_ receptor binding assays, inhibition of specific receptor binding was measured with cold ketanserin (100 nM, 10 µM). For 5-HT_2C_ receptor binding assays, inhibition of specific receptor binding was measured using cold SB 242,084 (100 nM, 10 µM). Inhibition of specific 5-HT receptor binding by the respective standards is summarized in Supplementary Table 6.

### Opioid receptor binding assays

#### Sample preparation

Receptor binding studies were conducted on CHO cells expressing µ-, κ-, or δ-opioid receptors. Rat µ-opioid receptor-expressing cells were provided by Dr. Thomas Murray (CHO-MOR) and human κ-, and δ-opioid receptor-expressing cells were provided by SRI International (CHO-KOR and CHO-DOR). CHO-MOR and CHO-KOR cell lines were maintained in DMEM supplemented with 10% FBS (Atlas Biologicals) and either 400 μg/ml or 200 μg/ml G418, respectively. CHO-DOR cells were maintained in DMEM supplemented with 10% FCS and 200 μg/mL G418. Upon reaching confluence, cells were harvested for membrane preparation. Cell membranes were prepared in 50 mM Tris buffer (pH 7.5 at 4°C). Cells were collected via scraping and centrifuged at 500 x *g* for 15 min. The cell pellet was homogenized in 2 mL buffer via polytron, diluted with 11 mL buffer, and centrifuged at 40000 x *g* for 15 min. The homogenized pellet was washed and recentrifuged, and finally resuspended at 3 mg protein/mL in buffer to determine protein content. Homogenates were stored at −70°C until use.

#### Binding assays

Binding assays were conducted using [^3^H]DAMGO (0.6 nM), [^3^H]U69,593 (0.8 nM), or [^3^H]DPDPE (0.5 nM), at the µ-, κ-, and δ-opioid receptors, respectively. The assay was performed in triplicate in a 96-well plate using 50 mM Tris buffer (pH 7.7, room temperature). Nonspecific binding was determined with 1 µM of unlabeled radioligand. Cell membranes were incubated with the appropriate radioligand and test compound at 25°C for 60 min. Incubation was terminated by rapid filtration through glass fiber filter paper presoaked in 0.1% polyethylenimine on a Tomtec cell harvester. Filters were dried and spotted with scintillation cocktail before counting for 2 min on a Perkin Elmer microBetaplate 1450 liquid scintillation counter. Nonspecific binding was determined with 1 µM of the unlabeled form of each radioligand.

#### Opioid receptor binding standards

Specificity of radioligand binding at the respective opioid receptors was measured via non-radiolabeled standards. For δ- and µ-opioid receptor binding assays, inhibition of specific receptor binding was measured using naltrexone (100 nM, 10 µM). For κ-opioid receptor binding assays, inhibition of specific receptor binding was measured with nor-BNI (100 nM, 10 µM). Inhibition of specific opioid receptor binding by the respective standards is summarized in Supplementary Table 7.

### Biogenic amine transporter binding assays

#### Sample preparation

Assay methods were adapted from previously described studies^59^. Briefly, HEK293 cells expressing either human dopamine transporter (HEK-hDAT) or human serotonin transporter (HEK-hSERT) were cultured in DMEM supplemented with 5% FBS, 5% BCS, 0.05 U penicillin/streptomycin, and 2 µg/mL puromycin. HEK293 cells expressing human norepinephrine transporter (HEK-hNET) were cultured in DMEM supplemented with 10% FBS, 0.05 U penicillin/streptomycin, and 300 μg/ml G418. Cells were grown to 80% confluence on 150 mm dishes. To prepare cell membranes, adherent cells were washed with Ca^2+^/Mg^2+^-free PBS, followed by addition of lysis buffer (10 mL; 2 mM HEPES with 1 mM EDTA). After 10 min, cells were scraped from plates and centrifuged at 30000 x *g* for 20 min. The pellet was resuspended in 0.32 M sucrose using a polytron homogenizer for 10 sec. The resuspension volume was dependent on the density of binding sites and was chosen to reflect binding of 10% or less of the total radioactivity.

#### Binding assays

Each assay tube contained 50 µL of membrane preparation (∼10-15 μg of protein), 50 µL of compound used to define non-specific binding, or buffer (Krebs-HEPES, pH 7.4; 122 mM NaCl, 2.5 mM CaCl_2_, 1.2 mM MgSO_4_, 10 μM pargyline, 100 μM tropolone, 0.2% glucose and 0.02% ascorbic acid, buffered with 25 mM HEPES), 25 µL of [^125^I]RTI-55 (40-80 pM final concentration) and additional buffer sufficient to bring the final volume to 250 µL. Membranes were preincubated with drug for 10 min prior to the addition of the [^125^I]RTI-55. Assay tubes were incubated at 25°C for 90 min. Binding was terminated by filtration over GF/C filters using a Tomtec 96-well cell harvester. Filters were washed for 6 sec with ice-cold saline. Scintillation fluid was added to each square and remaining filter radioactivity was determined using a Perkin Elmer μ-or beta-plate reader. Specific binding was defined as the difference in binding observed in the presence and absence of 5 μM mazindol (HEK-hDAT, HEK-hNET) or 5 μM imipramine (HEK-hSERT). Two independent competition experiments were conducted with triplicate determinations.

#### Biogenic amine transporter binding standard

Specificity of radioligand binding at the respective opioid receptors was measured via non-radiolabeled cocaine (100 nM, 10 µM). Inhibition of specific transporter binding is summarized in Supplementary Table 8.

### Off-target *in vitro* binding screen

A broad receptor and enzyme target binding screen to identify potential off-target BrMeI binding was performed commercially (Eurofins Cerep Panlabs, France) as described previously^60^. BrMeI was screened at 100 nM and 10 µM concentrations with each experiment conducted in duplicate. Membrane homogenates from stable cell lines expressing each receptor or enzyme (primarily human) were incubated with the respective radioligand in the presence of either BrMeI or reference control compounds. Respective reference compounds were tested concurrently with the test compound to assess assay reliability. Nonspecific binding for each target was ascertained in the presence of the specific agonist or antagonist. Following incubation, samples were vacuum-filtered through glass fiber filters, rinsed with ice-cold buffer using a 48-sample or 96-sample cell harvester, and filter radioactivity was measured via scintillation counter. Binding was calculated as % specific binding inhibition for each target.

### hERG channel activity assay

BrMeI was commercially evaluated for activity at the hERG (encoded by the human ether-a-go-go gene) K^+^ channel for potential cardiac toxicity (Eurofins Panlabs, Redmond, WA) similar to previous studies^61^. hERG K^+^ channel tail current was measured by the automated patch clamp (Qpatch 16) approach^62^. CHO-KI cells stably transfected with human hERG cDNA were used for the assay. Cells were harvested via trypsinization and maintained in serum-free medium at room temperature prior to recording. Cells were washed and resuspended into extracellular solution (137 mM NaCl, 4 mM KCl, 1.8 mM CaCl_2_, 1 mM MgCl_2_, 10 mM D(+)-glucose, 10 mM HEPES, pH 7.4). For recordings, the intracellular pipette solution consisted of 130 mM KCl, 10 mM NaCl, 1 mM MgCl_2_, 10 mM EGTA, 5 mM MgATP, 10 mM HEPES (pH 7.2). Upon achieving a whole cell configuration, cells were held at −80 mV. A 50 ms pulse to −40 mV was delivered to measure the leaking current which was subtracted from the tail current. Cells were subsequently depolarized to +20 mV for 2 sec, followed by a 1 sec pulse to −40 mV to induce tail currents. This recording procedure was repeated every 5 sec to monitor current amplitude. Tail current amplitudes were measured in response to different BrMeI concentrations (0.01-100 µM, in DMSO) applied sequentially. All experiments were performed at room temperature in duplicate. The reference compound control E-4031 was tested in parallel at multiple concentrations to obtain an IC_50_ curve. Data were analyzed to yield the % hERG channel inhibition by comparing tail current amplitude prior to and following BrMeI application; the current difference was normalized to control. Dose-response curves were fit via 3-parameter logistic regression of Log[drug] concentration to generate IC_50_ values for BrMeI-induced hERG channel inhibition.

### Mouse microsomal stability assay

Phase I metabolic stability assays were carried out in mouse liver microsomes as previously described^63^. Reactions were carried out with 100 mM potassium phosphate buffer, pH 7.4, in the presence of the NADPH regenerating system (1.3 mM NADPH, 3.3 mM glucose 6-phosphate, 3.3 mM MgCl_2_, 0.4 U/mL glucose-6-phosphate dehydrogenase, 50 µM sodium citrate). Reactions, conducted in triplicate, were initiated via addition of liver microsomes to the incubation mixture (1 µM compound final concentration; 0.2 mg/mL microsomes). Negative controls in the absence of NADPH system determined specific cofactor-free degradation. Compound disappearance was monitored via LC/MS/MS. Briefly, chromatographic analysis was performed using an Accela ultra high-performance system consisting of an analytical pump, and an autosampler coupled with a TSQ Vantage mass spectrometer (Thermo Fisher Scientific). Separation of the analyte from potentially interfering material was achieved at ambient temperature using an Agilent Eclipse Plus column (Agilent, Santa Clara, CA; 100 x 2.1mm i.d.) packed with a 1.8 µm C18 stationary phase. Mobile phase was composed of 0.1% formic acid in acetonitrile and 0.1% formic acid in water with gradient elution, starting with 10% (organic) linearly increasing to 99% up to 2.5 min, maintaining at 99% (2.5-3.5 min) and re-equilibrating to 10% by 4.5 min. The total run time for each analyte was 5.0 min.

### *In vivo* rodent studies

#### Animal husbandry

Animals were housed in accordance with NIH guidelines along with ARRIVE guidelines for reporting animal research^64, 65^. Mice and rats were housed in cages with a 12:12 light:dark cycle and had access to food and water *ad lib* at all times unless indicated otherwise. All efforts were made to ameliorate animal suffering.

#### Institutional approvals

All animal work was approved through the Institutional Animal Care and Use Committees at the University of Pittsburgh, Albert Einstein College of Medicine, Johns Hopkins University, and the University of Toronto.

#### Sex as a biological variable

Only male rodents were used to avoid the impact of estrus on hormone secretion and islet physiology. Additionally, our choice was based on recent work showing that male rodent β-cells possess transcriptomic signatures more similar to T2D compared to females^66, 67^.

#### *In vivo* pharmacokinetics

All pharmacokinetics assays were conducted in 6-8-week-old male CD1 mice (Harlan, Indianapolis, IN) weighing 20-25 g using similar methods to those described earlier^68–70^. BrMeI and unmodified bromocriptine were dosed at 10 mg/kg intravenously (i.v.) with dosing based on earlier metabolic studies in rodents^5, 71, 72^. Following administration, blood was collected via cardiac puncture at 15 min and 1 h post-dose (n=3 mice/time point). Plasma was harvested from blood by centrifugation at 3000 x *g*. Whole brains were collected, and all samples were stored at −80°C until analysis. Samples were extracted from the plasma using a one-step protein precipitation method with acetonitrile containing losartan (500 nM) as an internal standard. Extracts were vortexed and then centrifuged at 10000 x *g* (10 min, 4°C). For analysis of analytes in brain, whole brains were weighed and homogenized in 4 times the volume of acetonitrile (also containing internal standard losartan), followed by vortexing and centrifugation for sample extraction Once extracted, aliquots of the plasma and brain samples were diluted with water, transferred to a 250 µL polypropylene vial sealed with a Teflon cap, and analyzed via LC/MS/MS. Chromatographic analysis was performed on an Accela ultra high-performance system consisting of an analytical pump, and an autosampler coupled with TSQ Vantage mass spectrometer (see Supplementary Figure S5). Calibration standards were prepared from naïve mouse plasma or brain samples spiked with known concentrations of BrMeI and unmodified bromocriptine. Plasma concentrations (nmol/mL) and brain tissue concentrations (pmol/g) of the respective drugs were ascertained and plotted at the respective time points.

#### Diet

3-4-month-old wild-type C57BL/6J mice used in the glucose and insulin tolerance tests were fed a Western diet (5TJN: 40% fat, 44% carbohydrate, 16% protein; TestDiet) whose composition is analogous to previously described formulations^73, 74^. Animals were fed this Western diet for 12 weeks alongside drug administration.

#### Drug administration

Bromocriptine and BrMeI were dissolved in a solution of 50% DMSO in water (50% DMSO/50% dH_2_O) to a final concentration of 2 mg/mL and administered to mice i.p. Vehicle was defined as 50% DMSO/50% dH_2_O only. For glucose and insulin tolerance testing in mice, animals were administered the respective drugs or vehicle for 12 weeks via daily intraperitoneal (i.p.) administration. In a subset of glucose tolerance testing (GTT) studies, mice were administered either bromocriptine or vehicle daily via an intracerebroventricular (i.c.v.) route for 8 weeks prior to testing. At the 8-week time point, bromocriptine GTT data was compared between i.c.v. versus systemic bromocriptine administration. As a control, 8 weeks of systemic bromocriptine was sufficient to improve glucose tolerance in dysglycemic Western diet-fed mice (data not shown). All i.p. and i.c.v. treatments were conducted at the same time of day to control for potential circadian effects.

#### Intracerebral ventricular drug administration in mice

I.c.v. drug administration in adult mice was conducted as described earlier^75^. Mice were anesthetized with ketamine/xylazine for implantation of custom indwelling 3.5 mm guide cannulas (Plastics One Inc., Roanoke, VA) into the third cerebral ventricle using a stereotaxic apparatus (coordinates from bregma: anteroposterior, −0.3 mm; dorsoventral, −3 mm). Mice were allowed to recover for at least a week prior to i.c.v. drug administration.

#### Glucose tolerance testing

Oral glucose tolerance testing (OGTT) was conducted as previously described^75^. Briefly, Western diet-fed wild-type C57BL/6J on Western diet underwent a 6-h daytime fast (food away at 8AM, OGTT at 2PM) followed by a glucose challenge with a glucose load of 2 mg/kg (10% glucose stock). Glucose measurements were made from tail vein blood samples drawn at 0, 15-, 30-, 60- and 120-min following glucose challenge. Samples were measured immediately in duplicate using Contour glucometer sticks (Bayer Corp., Whippany, NJ).

#### Insulin tolerance testing

Western diet-fed wild-type C57BL/6J mice underwent a 6-h daytime fast, which was followed by intraperitoneal administration of 1.5 U/kg human insulin. Glucose values were determined from blood collected at 0, 15-, 30-, 60- and 120-min following insulin administration via tail bleed. Blood glucose samples were measured immediately in duplicate using Contour glucometer sticks. Insulin was measured in parallel via a quantitative ELISA kit (Linco Research Inc., St. Charles, MO).

#### Intracerebroventricular L-DOPA studies in rats

Male Sprague-Dawley rats underwent intracerebroventricular (i.c.v.; 3^rd^ ventricle) surgeries as described previously^20, 76^. A cannula was stereotaxically placed into the third cerebral ventricle under isoflurane anesthesia (coordinates from bregma: anteroposterior, −2.5 mm; mediolateral, 0.0 mm; dorsoventral, −8.0 mm). The cannula was secured in place with stainless steel screws (McCray Optical Supply Inc., Scarborough, ON, Canada) and dental cement, and kept patent with a steel stylet. Following approximately 1 week of recovery, rats underwent blood vessel cannulation for future blood sampling as earlier^20^. Rats were monitored closely after surgeries with behavior and body weight recorded daily to ensure successful recovery. Following ∼4 days of recovery from the last surgery, rats were food-restricted overnight and then infused (5 µL/h) i.c.v. with 10 mM L-DOPA or vehicle (saline) for 3.5 h. Acute potential changes in glycemic control were determined via periodic measurement of plasma glucose 0, 60-, 90-, 190-, 200-, and 210-min with a glucose analyzer (Analox Instruments, Stourbridge, UK). Plasma insulin was measured via ELISA (Mercodia, Winston Salem, NC).

### Statistics

Data are represented as means ± SEM for all experimental replicates. We used multi-factor analyses of variance (ANOVA; α = 0.05) to compare between-group differences. This included repeated measures two-way ANOVAs to compare between-group differences in glucose and insulin tolerance tests; Tukey’s multiple comparison tests were conducted for multiple post hoc comparisons. Statistical analyses were performed with GraphPad Prism. Significance was accepted at p < 0.05.

## Results

### Design and synthesis of quaternary MeI analogues of DA D_2_-like receptor-targeted drugs

We aimed to establish a family of pharmacological tools to selectively manipulate DA D_2_-like receptor signaling in the periphery while limiting their actions in the CNS. Based on earlier approaches employing MeI quaternization to produce peripherally-limited drugs^77–80^, we hypothesized that addition of a quaternary MeI group would similarly render D_2_-like receptor agonists less BBB-permeable. We first synthesized quaternary MeI salts of full D_2_-like receptor agonists that broadly target both D2R and D3R including (-)-quinpirole (**Figure 1A**) and bromocriptine (**Figure 1B**), producing MeI analogues AB01-59 and AB01-60, respectively. To focus more specifically on D2R, we quaternized sumanirole, a D2R-selective agonist, to generate MPE01-05 (**Figure 1C**). We also synthesized MeI analogues of D3R-preferential agonists (±)-7-OH-DPAT (AB01-62, **Figure 1D**) and (+)-PD128,907 (AB01-117, **Figure 1E**). To target peripheral D4R, we generated a quaternary MeI analogue of CAB03-015, a D4R-selective agonist^51^ (AB01-102, **Figure 1F**). Additionally, we synthesized an MeI analogue of aripiprazole which is a D_2_-like receptor partial agonist and is in clinical use as an APD^81, 82^ (AB01-61, **Figure 1G**). Finally, we generated an MeI analogue of a D_2_-like receptor antagonist by quaternizing L741,226 (MPE01-06, **Figure 1H**). Following synthesis, the respective MeI-modified compounds were evaluated to be >95% pure based on NMR, elemental analysis, HRMS (high resolution mass spectroscopy which agrees within ± 5 ppm) and MS/MS (ESI in positive mode) (**Supplementary Table 1**).

**Figure 1.**
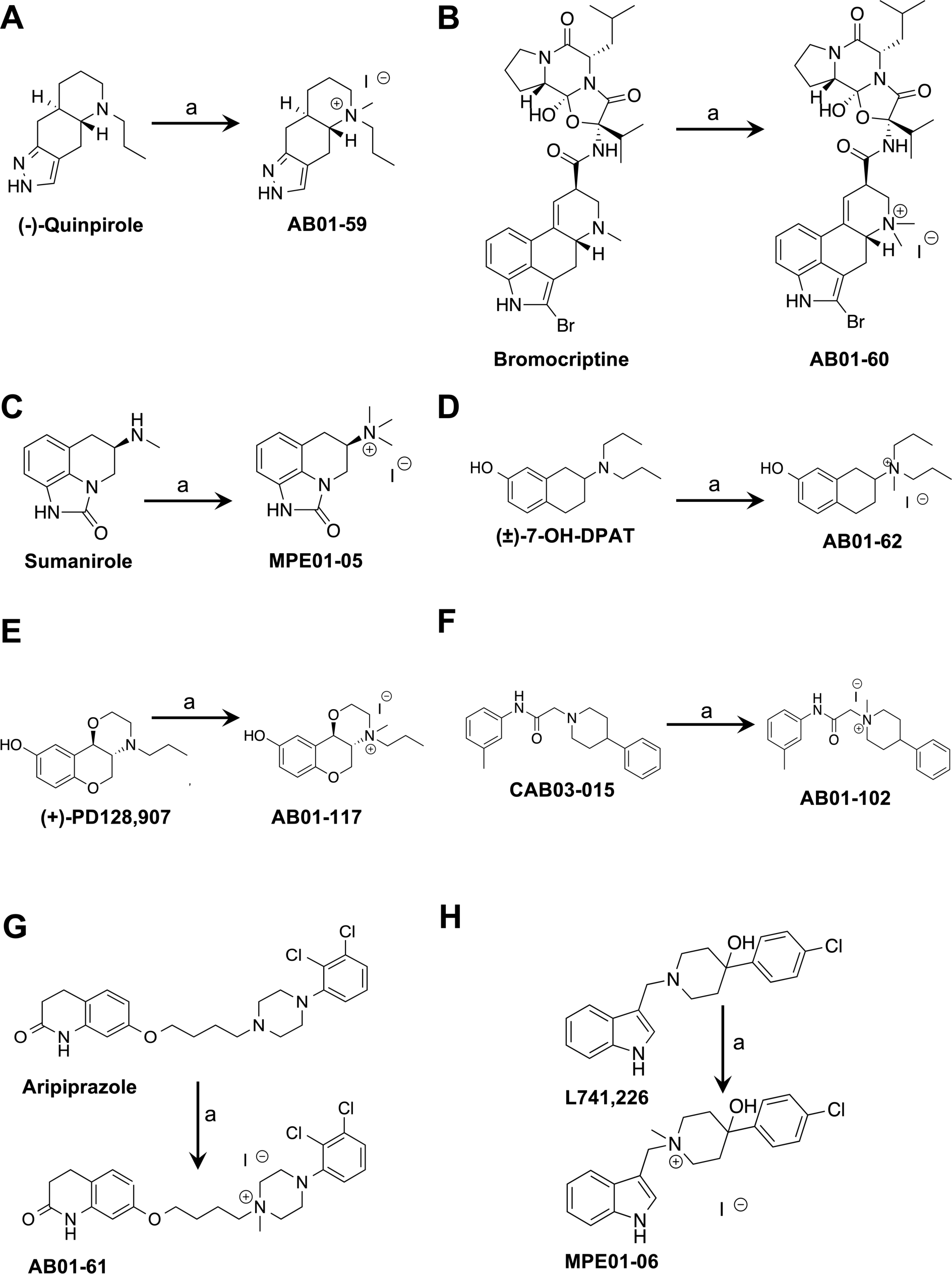
Drug design schemes of quaternary methiodide analogues of dopamine D_2_-like receptor-targeted drugs. To selectively manipulate dopamine D_2_-like receptor signaling in the periphery, drugs that target dopamine D_2_-like receptors were modified via addition of a quaternary methiodide (MeI) group to render them less permeable to the blood brain barrier. These MeI analogues are based on D_2_-like receptor agonists that preferentially target one or more of the receptors **(A-F)**: AB01-59 (**A**, (-)-quinpirole MeI), AB01-60 (**B**, bromocriptine MeI), MPE01-05 (**C**, sumanirole MeI), AB01-62 (**D**, (±)-7-OH-DPAT MeI), AB-01-117 (**E**, (+)-PD128,907 MeI), AB01-102 (**F**, CAB03-015 MeI); **(G)** a D_2_-like receptor partial agonist: AB01-61 (aripiprazole MeI); and **(H)** a D_2_-like receptor antagonist: MPE01-06 (L741,226 MeI). a = CH_3_I, acetone or EtOH, 24h.

### Characterization of D_2_-like receptor binding properties of MeI analogues

We ascertained whether MeI quaternization altered the D_2_-like receptor binding affinities of the MeI analogues by measuring dissociation constant (K_i_) values. Overall, the absolute K_i_ values were higher for all MeI analogues at human D2R and D3R compared to their respective parent compounds when in competition with agonist radioligand [^3^H]-(R)-(+)-7-OH-DPAT (**Table 1**). Nevertheless, some MeI analogues exhibited greater alterations in D2R and D3R binding affinities compared to others. For example, the MeI analogue of bromocriptine, AB01-60, continued to bind to D2R and D3R with relatively high affinity, demonstrating only 5.3-fold and 8.5-fold diminished affinities at D2R and D3R, respectively, compared to its unmodified parent compound. In contrast, the MeI analogue of (±)-7-OH-DPAT, AB01-62, showed 179-fold and 365-fold decreased binding affinity at D2R and D3R versus unmodified (±)-7-OH-DPAT. Just as importantly, MeI modification altered the relative balance of D2R versus D3R binding affinities for some of the analogues, while leaving others unchanged. AB01-59 showed preferential binding to D2R over D3R compared to unmodified (-)-quinpirole as indicated by a 6.3-fold-increase in the D3R/D2R ratio. In contrast, AB01-102, AB01-61, and MPE01-06 showed increased D3R affinity versus D2R (**Table 1**), suggesting that local changes in polarity following MeI addition may play a key role in determining D2R/D3R receptor binding properties. Finally, we measured the binding affinities of D4R-selective agonist CAB03-015 and its MeI analogue AB01-102 at human D4R, finding a 61.7-fold loss of binding affinity for AB01-102 compared to its unmodified parent (**Supplementary Table 2**).

**Table 1.** Radioligand competition D_2_-like receptor binding data. Equilibrium dissociation constants (K_i_) indicate human D2R and D3R binding affinities for dopamine D_2_-like receptor-selective drugs and their respective quaternary MeI analogues. The ratio of D3R/D2R binding affinity was calculated alongside each compound. All compounds were tested with [^3^H]-(R)-(+)-7-OH-DPAT, except for L741,626 and MPE01-06 which employed [^3^H]N-methylspiperone. K_i_ values are represented as mean ± SEM with n ≥ 3 independent experiments performed in triplicate. ^a^Data previously reported in Bonifazi et al. (2017).

| Compound | D2R | D3R | D3R/D2R |
| --- | --- | --- | --- |
| | $K_i \pm \text{SEM (nM)}$ | $K_i \pm \text{SEM (nM)}$ | |
| **(-)-Quinpirole<sup>a</sup>** | 13.1 ± 2.2 | 6.76 ± 1.2 | 0.52 |
| **AB01-59** | 141.0 ± 52.4 | 459.0 ± 80.5 | 3.26 |
| **Bromocriptine** | 0.59 ± 0.17 | 2.05 ± 0.56 | 3.46 |
| **AB01-60** | 3.14 ± 0.34 | 17.50 ± 4.90 | 5.57 |
| **Sumanitrole<sup>a</sup>** | 46.3 ± 2.6 | 573 ± 118 | 12.38 |
| **MPE01-05** | 129.0 ± 20.5 | 1840 ± 271 | 14.26 |
| **(±)-7-OH-DPAT** | 5.09 ± 1.21 | 3.42 ± 0.72 | 0.67 |
| **AB01-62** | 912 ± 117 | 1250 ± 183 | 1.37 |
| **(+)-PD128,907** | 18.8 ± 4.7 | 2.65 ± 0.14 | 0.14 |
| **AB01-117** | 94.1 ± 20.5 | 18.7 ± 4.3 | 0.20 |
| **CAB03-015** | 140.0 ± 39.6 | 806 ± 145 | 5.76 |
| **AB01-102** | 1750 ± 276 | 2900 ± 456 | 1.66 |
| **Aripiprazole<sup>a</sup>** | 0.48 ± 0.05 | 3.57 ± 0.79 | 7.44 |
| **AB01-61** | 437 ± 60 | 1970 ± 360 | 4.51 |
| **L741,626** | 8.47 ± 0.99 | 54.1 ± 13.0 | 6.39 |
| **MPE01-06** | 1660 ± 327 | 1130 ± 440 | 0.68 |

### Functional characterization of MeI analogues

We evaluated the activity of our MeI analogues via the mitogenesis functional assay, where we measured D_2_-like receptor-mediated increases in the incorporation of [^3^H]thymidine in CHO cells expressing either human D2R or D3R^53^ (**Figure 2**). The canonical D2R/D3R agonists (-)-quinpirole and bromocriptine and their respective MeI analogues produced robust concentration-dependent D2R- and D3R-mediated increases in [^3^H]thymidine incorporation (**Figures 2A-2D**). However, while the efficacies of both AB01-59 and AB01-60 were largely unchanged compared to their respective unmodified parents, AB01-59 displayed diminished potency in inducing D2R- and D3R-mediated mitogenesis (24.9-fold, D2R; 14.4-fold, D3R) compared to AB01-60 (15.6-fold, D2R; 11.2-fold, D3R) (**Table 2**). Notably, MPE01-05, the MeI analogue of sumanirole, did not drive significant changes in either efficacy or potency in D2R- and D3R-mediated mitogenesis (**Figures 2E, 2F**). In contrast, AB01-62 and AB01-117, MeI analogues of preferential D3R agonists, demonstrated substantial loss of potency in D3R-mediated mitogenesis (AB01-62: 1107.4-fold decrease; AB01-117: 60.3-fold decrease) despite retaining their efficacies (**Figures 2G, 2H, Table 2**). Consistent with aripiprazole’s partial agonism of D2R and D3R, MeI analogue AB01-61 also displayed partial agonist properties in D2R- and D3R-mediated mitogenesis with near-identical efficacies. However, AB01-61 showed markedly reduced potencies compared to aripiprazole (D2R, 1605.3-fold decrease; D3R, 90.7-fold decrease), suggesting that this analogue has little functional activity at either D2R or D3R (**Figures 2I**, 2J, **Table 2**). As expected, MPE01-06, the MeI analogue of D2R/D3R antagonist L741,626, had no significant functional effects on mitogenesis in our assay (<1% of the maximum (-)-quinpirole response; **Figures 2K**, 2L, **Table 2**). Lastly, since CAB03-015 is a D4R-selective agonist, we tested the activity of the drug and its MeI analogue AB01-102 in a D4R-mediated adenylate cyclase activity/cyclic AMP (cAMP) formation assay. Both drugs behaved as partial agonists, exhibiting similar efficacies in eliciting D4R-driven decreases in forskolin-stimulated adenylate activity and the resulting cAMP formation. However, AB01-102 showed strongly diminished potency (731.9-fold decrease) compared to unmodified CAB-03-015 (**Supplementary Table 3**).

**Figure 2.**
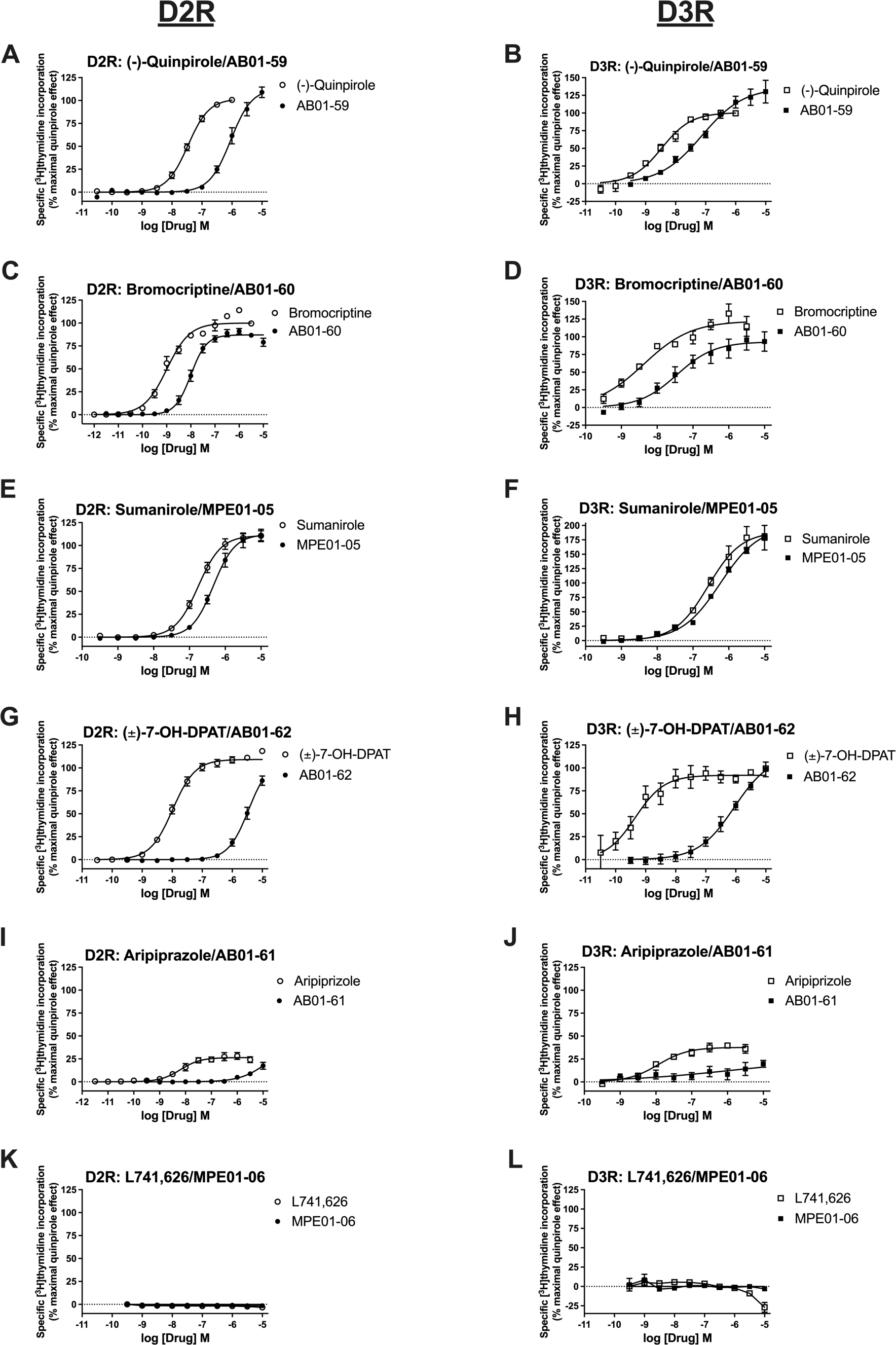
Impact of quaternary methiodide analogues of dopamine D_2_-like receptor-targeted drugs on D2R- and D3R-mediated mitogenesis. Dose response curves demonstrating the impact of dopamine D_2_-like receptor-selective drugs (open circles or squares) and their quaternary methiodide (MeI) analogues (solid circles or squares) on mitogenesis in cultured CHO cells expressing human D2R **(A, C, E, G, I, K)** or human D3R **(B, D, F, H, J, L)**. Mitogenesis was assessed according to [^3^H]thymidine incorporation. **(A-H)** All canonical D2R/D3R agonists including (-)-quinpirole **(A, B)**, bromocriptine **(C, D)**, sumanirole **(E, F)**, (±)-7-OH-DPAT **(G, H)**, and their respective MeI analogues showed dose-dependent stimulation of D2R- and D3R-mediated mitogenesis. Compared to their unmodified parent drugs, MeI-modified agonist analogues possessed similar efficacies but diminished potencies. **(I, J)** D2R/D3R partial agonist aripiprazole exhibited decreased efficacy for both D2R- and D3R-mediated mitogenesis stimulation compared to full agonists (**A-H**). MeI analogue AB01-61 showed almost total loss of efficacy and potency versus its unmodified parent compound. **(K, L)** D2R/D3R antagonist L741,626 and its MeI analogue MPE01-06 had no effect on D2R- and D3R-mediated mitogenesis. Results for [^3^H]thymidine incorporation for each drug were normalized to % maximal quinpirole effect. Data are represented as means ± SEM for all experimental replicates and were performed in triplicate from n ≥ 3 independent experiments.

**Table 2.** Functional activity in D2R- and D3R-mediated mitogenesis assays. EC_50_ and *E_max_* values were calculated from dose-response curves of drug-stimulated mitogenesis in CHO cells expressing either human D2R or D3R. EC_50_ values are represented as the mean ± SEM. E_max_ values represent % maximal efficacy relative to quinpirole stimulation. Data are from n ≥ 3 independent experiments performed in triplicate. ND, not determined.

| Compound | D2R |  | D3R |  |
| --- | --- | --- | --- | --- |
|  | EC<sub>50</sub> ± SEM (nM) | E<sub>max</sub> | EC<sub>50</sub> ± SEM (nM) | E<sub>max</sub> |
| (-)-Quinpirole | 35.8 ± 5.8 | 102% | 3.9 ± 1.0 | 102% |
| AB01-59 | 890 ± 190 | 109% | 56 ± 14 | 124% |
| Bromocriptine | 0.63 ± 0.21 | 99% | 3.4 ± 1.1 | 116% |
| AB01-60 | 9.8 ± 2.9 | 88% | 38 ± 11 | 91% |
| Sumanitrole | 164 ± 54 | 110% | 224 ± 24 | 180% |
| MPE01-05 | 472 ± 79 | 105% | 386 ± 76 | 172% |
| (±)-7-OH-DPAT | 10.8 ± 1.6 | 106% | 0.54 ± 0.17 | 97% |
| AB01-62 | 3520 ± 610 | 105% | 598 ± 58 | 102% |
| (+)-PD128,907 | ND |  | 0.78 ± 0.28 | 98% |
| AB01-117 | ND |  | 47 ± 14 | 98% |
| CAB03-015 | ND |  | >10000 (IC<sub>50</sub> = 2770 ± 310) | <1% (I<sub>max</sub> = 83%) |
| AB01-102 | ND |  | >8300 (IC<sub>50</sub> = >8500) | 6% (I<sub>max</sub> = 64%) |
| Aripiprazole | 3.8 ± 1.5 | 25% | 18.3 ± 5.6 | 29% |
| AB01-61 | 6100 ± 1300 | 29% | 1660 ± 670 | 27% |
| L741,626 | >10000 | <1% | >10000 | <5% |
| MPE01-06 | >10000 | <0% | >10000 | <2% |

### MeI analogues modify glucose-stimulated insulin secretion

We and our colleagues previously showed that human and rodent insulin-secreting pancreatic β-cells expressed D_2_-like receptors including D2R and D3R, and that agonist stimulation of these receptors by (-)-quinpirole and bromocriptine decreased glucose-stimulated insulin secretion (GSIS)^9, 12, 13, 27–29, 57, 83, 84^. Therefore, we further functionally tested the ability of several of the MeI analogues to directly signal through β-cell D_2_-like receptors by examining their impact on GSIS in INS-1E cells, an insulin-secreting rat pancreatic β-cell line^55^. As expected, unmodified (-)-quinpirole and bromocriptine dose-dependently reduced GSIS [(-)-quinpirole IC_50_ = 1.18 μM; bromocriptine IC_50_ = 0.18 μM] (**Figures 3A, 3B, Table 3**). AB01-59, the MeI analogue of (-)-quinpirole, similarly decreased GSIS, but with reduced efficacy (48.4% decrease) and potency (IC_50_ = 6.94 μM, 5.9-fold decrease) versus unmodified (-)-quinpirole (**Figure 3A**, **Table 3**). By comparison, bromocriptine MeI analogue AB01-60 functioned as a full agonist, completely retaining its efficacy, albeit with decreased potency (IC_50_ = 10.17 μM, 57.7-fold decrease) compared to unmodified bromocriptine (**Figure 3B**, **Table 3**). Interestingly, the D2R-preferential agonist sumanirole^85^ or D3R-preferential PD128,907^86^ and their respective MeI analogs exhibited markedly less efficacy and potency in GSIS inhibition than non-selective D2R/D3R agonists (-)-quinpirole or bromocriptine (**Figures 3C, 3D, Table 3**). This is consistent with our recent findings that both D2R and D3R must work in combination to effectively modulate GSIS^11^. Together, given AB01-60’s preservation of D2R/D3R binding affinity and efficacy in both GSIS and mitogenesis assays (**Tables 2, 3**), we chose this compound (henceforth termed bromocriptine MeI or BrMeI) as our lead compound.

**Figure 3.**
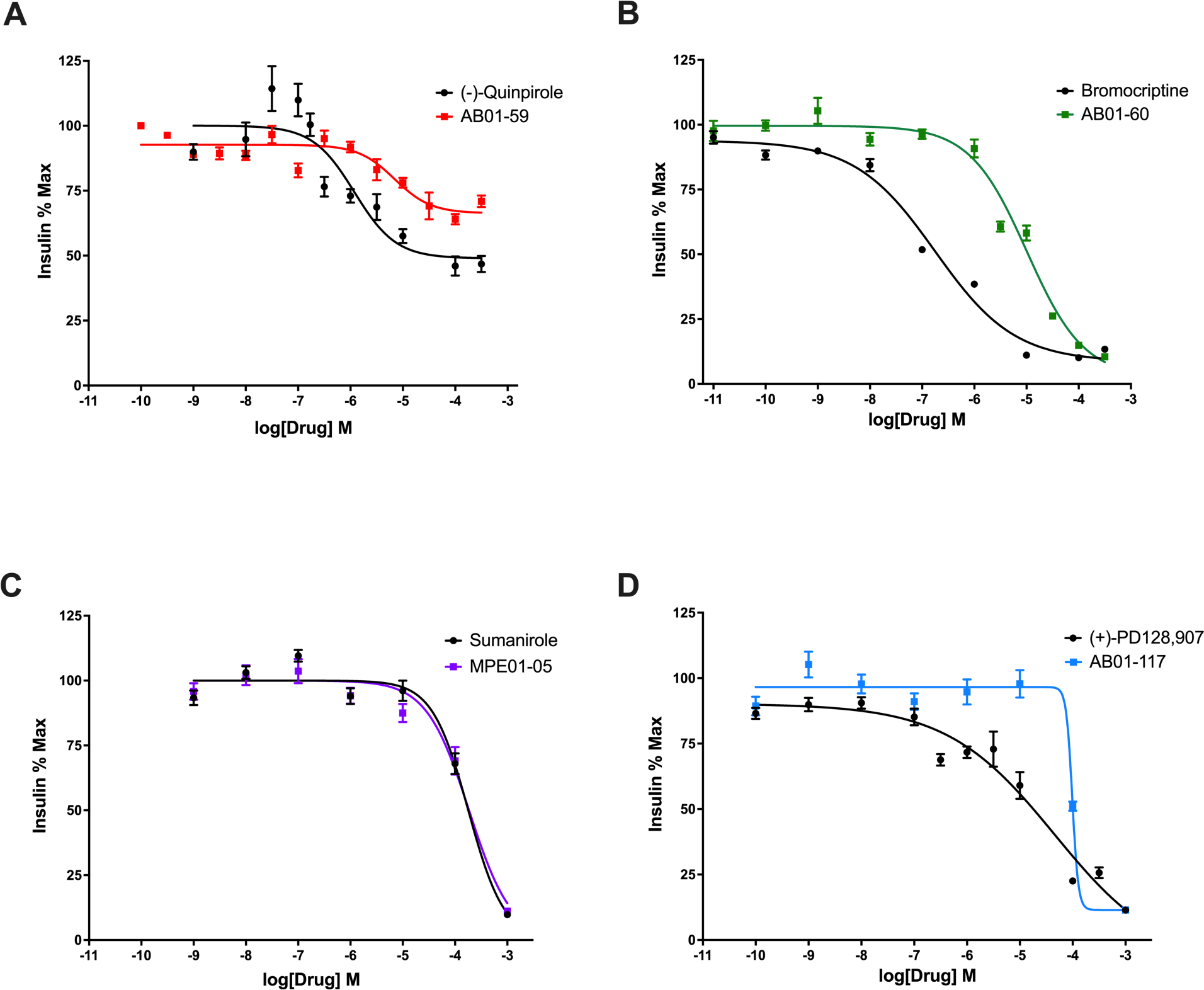
Impact of quaternary methiodide analogues on glucose-stimulated insulin secretion. Dose response curves demonstrating the impact of dopamine D_2_-like receptor-selective drugs (circles) and their quaternary methiodide (MeI) analogues (squares) on glucose-stimulated insulin secretion (GSIS) in INS-1E cells. **(A)** Treatment with D2R/D3R agonist (-)-quinpirole produced a dose-dependent reduction of GSIS (IC_50_ = 1.18 μM). (-)-Quinpirole MeI analogue AB01-59 also decreased GSIS with reduced efficacy (48.4% decrease) and potency (IC_50_ = 6.94 μM). **(B)** Bromocriptine was more potent than (-)-quinpirole in reducing GSIS (IC_50_=0.18 μM). Bromocriptine MeI analogue AB01-60 retained its efficacy but demonstrated decreased potency (IC_50_ = 10.17 μM) versus unmodified bromocriptine. **(C)** Selective D2R agonist sumanirole dose-dependently decreased GSIS with lower efficacy and potency (IC_50_=17.92 μM) than (-)-quinpirole or bromocriptine. Sumanirole MeI analogue MPE01-05 possessed nearly the same efficacy and similar potency (IC_50_=18.50 μM) as unmodified sumanirole. **(D)** Selective D3R agonist (+)-PD128,907 produced less potent (IC_50_ = 45.43 μM) GSIS reduction compared to drugs acting via selective D2R agonism or joint D2R/D3R agonism. (+)-PD128,907 MeI analogue AB01-117 was less potent (IC_50_ = 98.19 μM) but similarly efficacious as its unmodified parent. Insulin data were normalized to % maximal secreted insulin. Data are represented as means for all experimental replicates ± SEM and were from n ≥ 2 independent experiments.

**Table 3.**
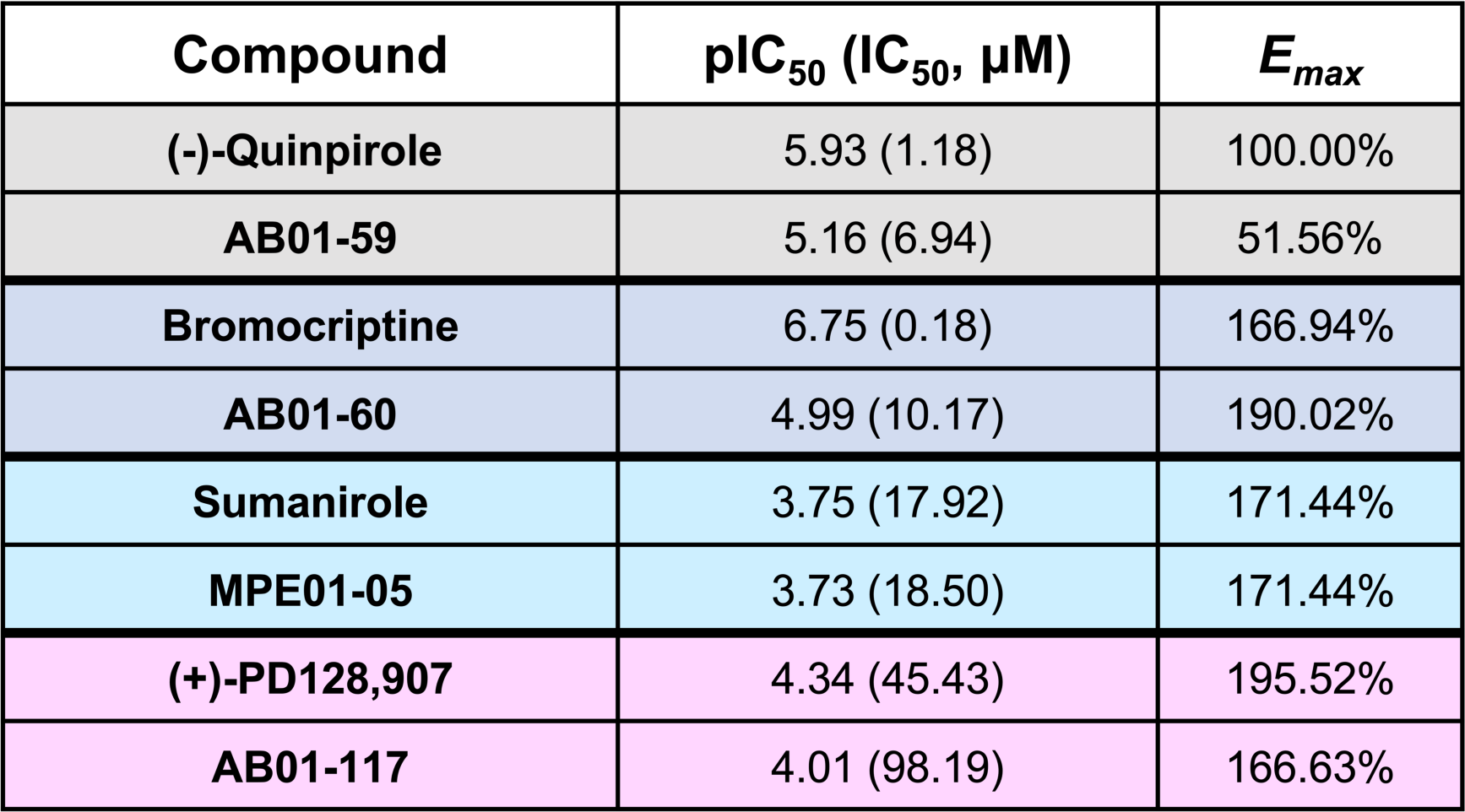
Drug potencies and efficacies on glucose-stimulated insulin secretion (GSIS). Potency data are represented by pIC_50_ with corresponding IC_50_ values in parentheses (in μM). *E_max_* values represent % maximal efficacy relative to (-)-quinpirole treatment.

### BrMeI preferentially recruits β-arrestin2 to D2R

We next characterized BrMeI-stimulated signaling through D2R. We focused on BrMeI’s ability to initiate D2R-mediated recruitment of intracellular effectors such as G protein Gα_i1_ and β-arrestin2, given their important metabolic roles in the regulation of hormone secretion^13, 32, 87–89^. We employed highly sensitive nanoBRET technology to determine if BrMeI-stimulated recruitment of Gα_i1_ and β-arrestin2 to D2R differed from that of either unmodified bromocriptine or endogenous ligand DA. In our assay, D2R was HaloTag-labeled with a bright fluorescent dye and either Gα_i1_ or β-arrestin2 was tagged with nanoluciferase (NanoLuc). In response to receptor recruitment, the resulting proximity of the receptor-effector pair enables luminescence from the NanoLuc-tagged effector to excite the receptor-bound dye, generating quantifiable fluorescence^90^. D2R stimulation by DA caused dose-dependent recruitment of Gα_i1_ (EC_50_ = 288.67 nM). Bromocriptine was much more potent than DA at eliciting Gα_i1_ recruitment to D2R (EC_50_ = 15.07 nM), albeit with 47% less efficacy. This suggested that bromocriptine functioned as a partial agonist in eliciting Gα_i1_ recruitment to D2R. BrMeI’s efficacy resembled unmodified bromocriptine, though with less potency (EC_50_ = 1514.71 nM) (**Figure 4A, Supplementary Table 4**). Importantly, we discovered that BrMeI more potently elicited β-arrestin2 recruitment to D2R (EC_50_ = 8.37 nM) compared to bromocriptine (EC_50_ = 20.13 nM) and DA (EC_50_ = 27.63 μM), while still retaining bromocriptine’s reduced efficacy (**Figure 4B, Supplementary Table 4**). These data suggest that BrMeI is an agonist of D2R that preferentially recruits β-arrestin2 over G proteins to D2R.

**Figure 4.**
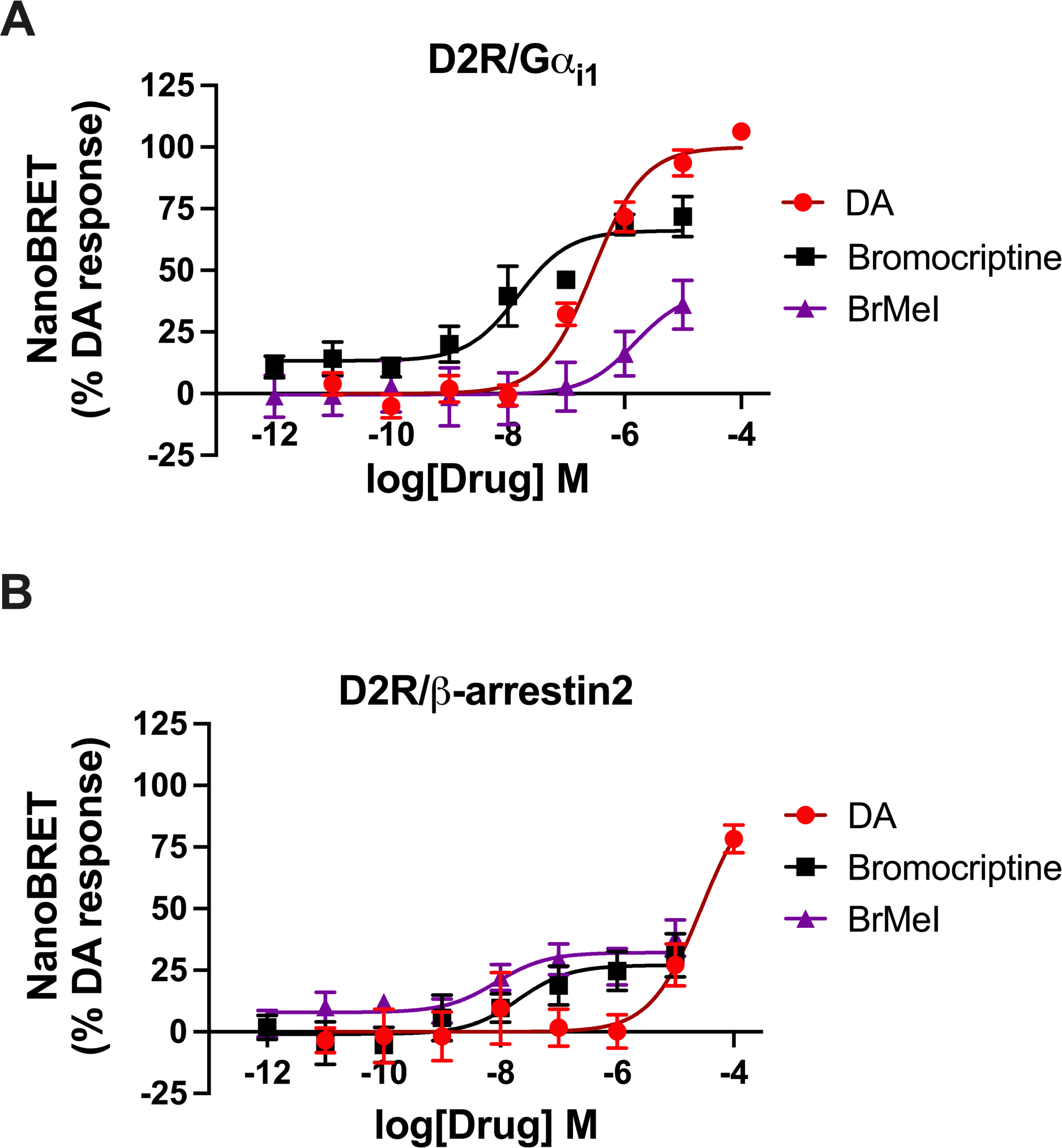
Bromocriptine methiodide stimulation causes preferential recruitment of β-arrestin2 to D2R. Concentration-response nanoBRET assays examining recruitment of G protein Gα_i1_ and β-arrestin2 to D2R in response to stimulation by DA (red circle), unmodified bromocriptine (black square), or bromocriptine methiodide (BrMeI, purple triangle). HEK-293T cells expressed HaloTag-labeled D2R and either NanoLuc-labeled Gα_i1_ or **β**-arrestin2 as the respective nanoBRET pairs. **(A)** Stimulation by DA, bromocriptine, or BrMeI resulted in dose-dependent Gα_i1_ recruitment to D2R. BrMeI was less efficacious or potent (EC_50_ = 1.52 μM) than either canonical D2R ligand DA (EC_50_=0.29 μM) or bromocriptine (EC_50_ = 15.07 nM). **(B)** BrMeI was more potent in eliciting β-arrestin2 recruitment to D2R (EC_50_ = 8.37 nM) compared to bromocriptine (EC_50_ = 20.13 nM) or DA (EC_50_ = 27.63 μM). BrMeI exhibited β-arrestin2 recruitment efficacy similar to bromocriptine but was 4-fold less efficacious than DA. NanoBRET data were baseline-corrected and normalized to % maximal DA response. Assays were performed in triplicate from n ≥ 3 independent experiments. Data are represented as mean ± SEM for all experimental replicates.

### *In vitro* target binding profile of BrMeI

We further characterized BrMeI’s binding specificity in a series of *in vitro* radioligand binding competition assays via an initial screen for binding to several families of G protein-coupled receptors (*e.g.*, DA, serotonin, opioid receptors) and biogenic amine transporters. Radioligand binding was successfully validated using established standards (**Supplementary Tables 5-8**). At submicromolar concentration (100 nM), BrMeI did not exhibit significant screen hits (**Supplementary Figure 1**). However, at a higher concentration (10 µM), BrMeI demonstrated strong D2R and D3R binding (92.2% and 99.0% inhibition of specific radioligand binding, respectively), consistent with its potencies in the GSIS and mitogenesis assays. In contrast, BrMeI did not significantly bind to D1R or D4R (**Supplementary Figure 1**). We additionally observed significant BrMeI binding to serotonin 5-HT_1A_, 5-HT_2A_, and 5-HT_2C_ receptors as well as to δ-, κ-, and µ-opioid receptors at the 10 µM concentration, which is in line with bromocriptine’s ability to similarly bind other receptors^41^. We found no biogenic amine transporter binding at either tested BrMeI concentrations (**Supplementary Figure 1**).

We next conducted a larger screen to detect potential off-target binding by BrMeI across a broad range of human receptors, channels, and enzymes. As above, we found no significant hits at the lower submicromolar 100 nM BrMeI concentration (**Supplementary Figure 2, Supplementary Figure 3**). At the higher 10 µM concentration, we identified BrMeI binding to ⍺_2_ adrenergic receptors, consistent with work showing bromocriptine’s ability to signal via both D_2_-like and ⍺_2_ adrenergic receptors^12, 91, 92^. At 10 μM BrMeI, we also found positive hits at A, ⍺, ⍺, D R, κ-_2+ +_ opioid, µ-opioid, M, M_2_, NK_1_, and NK_2_ receptors, L-type Ca^2+^ (diltiazem) and Na (site 2) channels as well as acetylcholinesterase (**Supplementary Figure 2, Supplementary Figure 3**). Finally, we tested BrMeI for hERG channel activity, finding concentration-dependent hERG tail current inhibition (IC_50_ = 8.65 µM) (**Supplementary Figure 4**), consistent with similar hERG activity by unmodified bromocriptine^93^.

### BrMeI undergoes Phase I metabolism in mouse microsomes identically to bromocriptine

We tested BrMeI for Phase I metabolic stability using mouse liver microsomes fortified with NADPH, comparing its stability to unmodified bromocriptine; fractions of drug remaining over time were measured via LC/MS/MS^70^. Incubation of BrMeI and bromocriptine in the presence of NADPH led to the complete disappearance of bromocriptine within 30 min (**Figure 5A**). Similarly, we found almost complete BrMeI loss within 30 min and total loss by 60 min (**Figure 5B**). These results indicate that BrMeI undergoes significant hepatic metabolism in a manner identical to unmodified bromocriptine.

**Figure 5.**
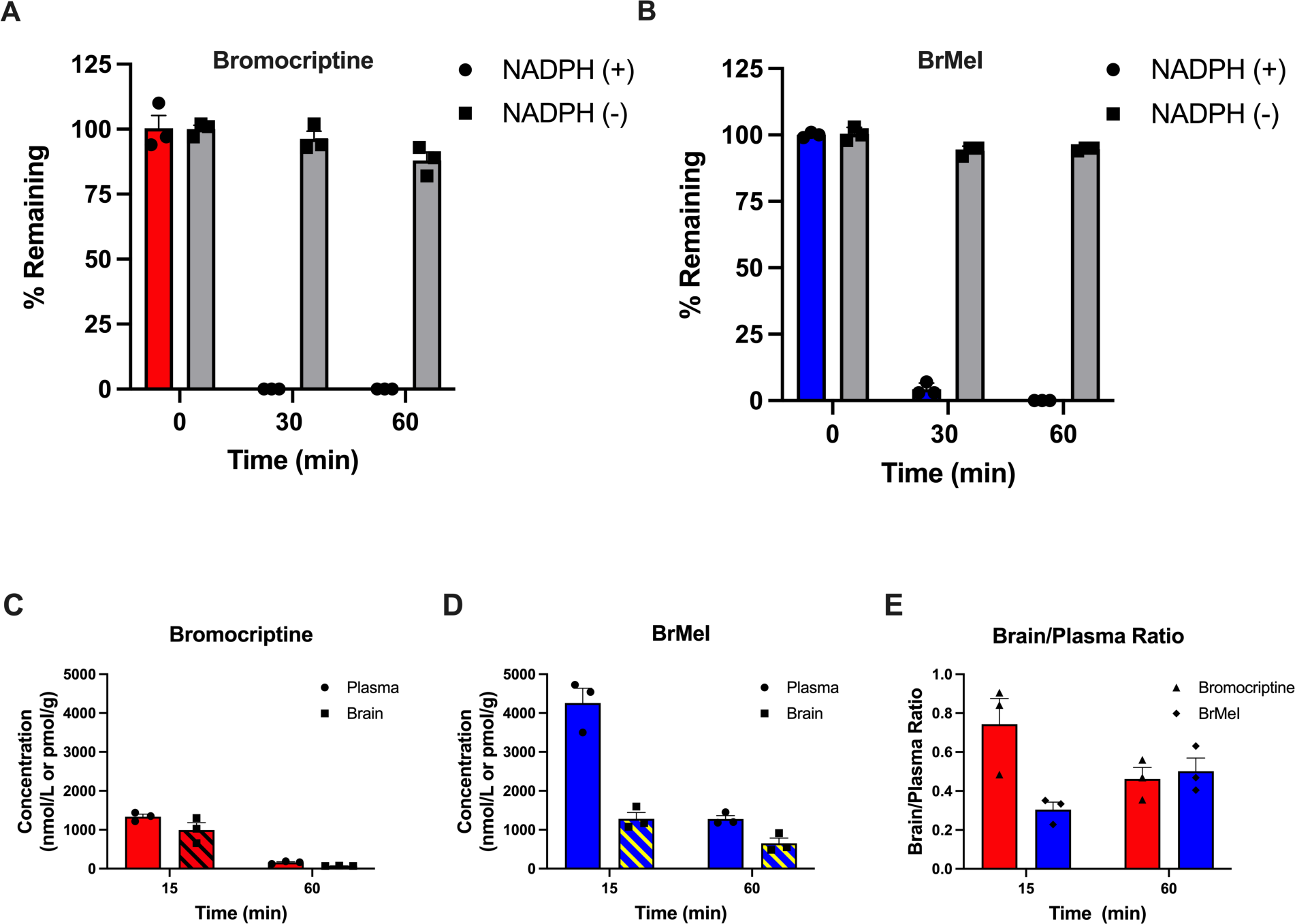
Phase I metabolic stability and pharmacokinetic analysis of BrMeI versus bromocriptine. **(A, B)** Phase I metabolic stability assay of BrMeI versus bromocriptine. BrMeI and bromocriptine (1 µM) were incubated with mouse liver microsomes in the presence or absence of NADPH (negative control). % drug remaining across time was measured by LC/MS/MS. Bromocriptine **(A)** and BrMeI **(B)** underwent significant Phase I metabolism in mouse liver microsomes. Bromocriptine was entirely metabolized within 30 min. For BrMeI, <5% of the drug similarly remained by 30 min and 0% within 60 min. Data are represented as the mean % remaining drug ± SEM (n = 3 per time point). **(C-E)** Pharmacokinetic analysis of BrMeI versus bromocriptine in mice. Brain and plasma concentrations of BrMeI and bromocriptine were measured at 15 min and 60 min following intravenous administration (10 mg/kg for both drugs). Bromocriptine **(C)** exhibited lower plasma levels at both time points compared to BrMeI **(D)**. Plasma levels of BrMeI remained 8-fold higher at 1 h compared to bromocriptine. **(E)** The brain-to-plasma ratios for BrMeI were lower at 15 min but similar at 60 min. Data are represented as the mean ± SEM (n=3 per group).

### Pharmacokinetic evaluation of BrMeI *in vivo*

We evaluated the pharmacokinetics of BrMeI versus unmodified bromocriptine *in vivo* in wild-type CD1 mice. Mice were administered 10 mg/kg BrMeI or bromocriptine via an intravenous (i.v.) route followed by LC/MS/MS measurements of plasma and brain drug levels 15- and 60-min post-administration (**Figure 5C-E, Supplementary Figure 5, Table 4**). At 15 min, the brain/plasma ratio for BrMeI was 2.5-fold lower compared to bromocriptine, suggesting that, acutely, BrMeI had lower brain penetrance and was preferentially localized to the periphery (**Figures 5C, 5D**). However, by 60 min, the brain/plasma ratios of BrMeI and bromocriptine were similar (bromocriptine, 0.45; BrMeI, 0.51), suggesting that BrMeI was not completely restricted from the CNS (**Figure 5E**). Rather, quaternary MeI modification significantly slowed BBB penetration of the drug. Importantly, by 60 min post-administration, most unmodified bromocriptine was cleared from both brain (92.9% decrease) and plasma (88.2% decrease) *in vivo* (**Table 4**). In contrast, significantly higher plasma and brain levels of BrMeI still remained at this later timepoint (**Figure 5**). Indeed, plasma BrMeI levels were ∼8-fold greater compared to bromocriptine at 60 min, suggesting that BrMeI possessed an *in vivo* pharmacokinetic profile distinct from unmodified bromocriptine which enabled increased duration of action.

**Table 4.** Quantification of *in vivo* pharmacokinetics of BrMeI versus bromocriptine in mice. Data are represented as the mean ± SEM (n=3 per group).

| Drug | Time (min) | Plasma Conc. (nmol/L) | Brain Conc. (pmol/g) | Brain/Plasma Ratio |
| --- | --- | --- | --- | --- |
| Bromocriptine | 15 min | 1336.7 ± 52.2 | 995.2 ± 188.0 | 0.74 ± 0.11 |
|  | 60 min | 158.6 ± 15.0 | 71.1 ± 2.3 | 0.45 ± 0.05 |
| BrMel | 15 min | 4259.1 ± 381.3 | 1282.3 ± 130.7 | 0.30 ± 0.03 |
|  | 60 min | 1277.1 ± 71.1 | 651.9 ± 133.7 | 0.51 ± 0.06 |

### Actions at both CNS and peripheral targets are required to effectively treat dysglycemia *in vivo*

Finally, we examined the metabolic impacts of BrMeI versus bromocriptine on systemic glucose homeostasis *in vivo* in a diet-induced obesity (DIO) model of T2D where wild-type C57BL/6J mice were chronically maintained on a Western diet to induce dysglycemia and obesity. As expected, vehicle-treated control mice exhibited fasting hyperglycemia and impaired glucose tolerance during oral glucose tolerance testing (OGTT). We discovered that systemic administration of bromocriptine produced significant improvements in glucose tolerance as indicated by a significant effect of drug [F(2, 29) = 13.10, p < 0.0001] as well as an interaction of drug × time [F(8, 116) = 4.178, p = 0.0002]. In contrast, systemic administration of BrMeI did not significantly modify glucose tolerance compared to vehicle (p > 0.05) (**Figure 6A**). Systemic drug treatment also lowered fasting blood glucose [F(2, 29) = 6.994, p = 0.0033]. While bromocriptine-treated mice exhibited significantly lower fasting glucose, BrMeI treatment had no significant effect (p > 0.05) (**Figure 6B**). Insulin tolerance testing (ITT) similarly revealed a significant effect of drug treatment on blood glucose [F(2, 21) = 6.761, p = 0.0054] driven by bromocriptine-induced improvements in insulin sensitivity (**Figure 6C**). Notably, systemic BrMeI’s effects on blood glucose during the ITT were intermediate to those of bromocriptine and vehicle. BrMeI significantly reduced blood glucose versus vehicle at the 60 min time point (p = 0.022, **Figure 6C**), suggesting that the drug may have partial efficacy on improving insulin sensitivity *in vivo*. We also measured the impact of systemic bromocriptine and BrMeI treatment on blood insulin during the initial 15 min of the ITT time course. There was a significant drug × time interaction [F(4, 42) = 10.57, p < 0.0001] on blood insulin driven by bromocriptine’s ability to significantly enhance the levels of circulating plasma insulin within 5 min of insulin administration versus vehicle (p = 0.0002) or BrMeI (p = 0.0003); BrMeI treatment did not significantly alter plasma insulin (p > 0.05) (**Figure 6D**). Together, these results were consistent with bromocriptine’s established role as a treatment for dysglycemia in T2D^42, 94, 95^. Interestingly, neither systemic bromocriptine nor BrMeI treatments had significant effects on weight at 12 weeks (p > 0.05) (**Figure 6E**), suggesting that drug-induced effects on dysglycemia could be dissociated from effects on weight gain.

**Figure 6.**
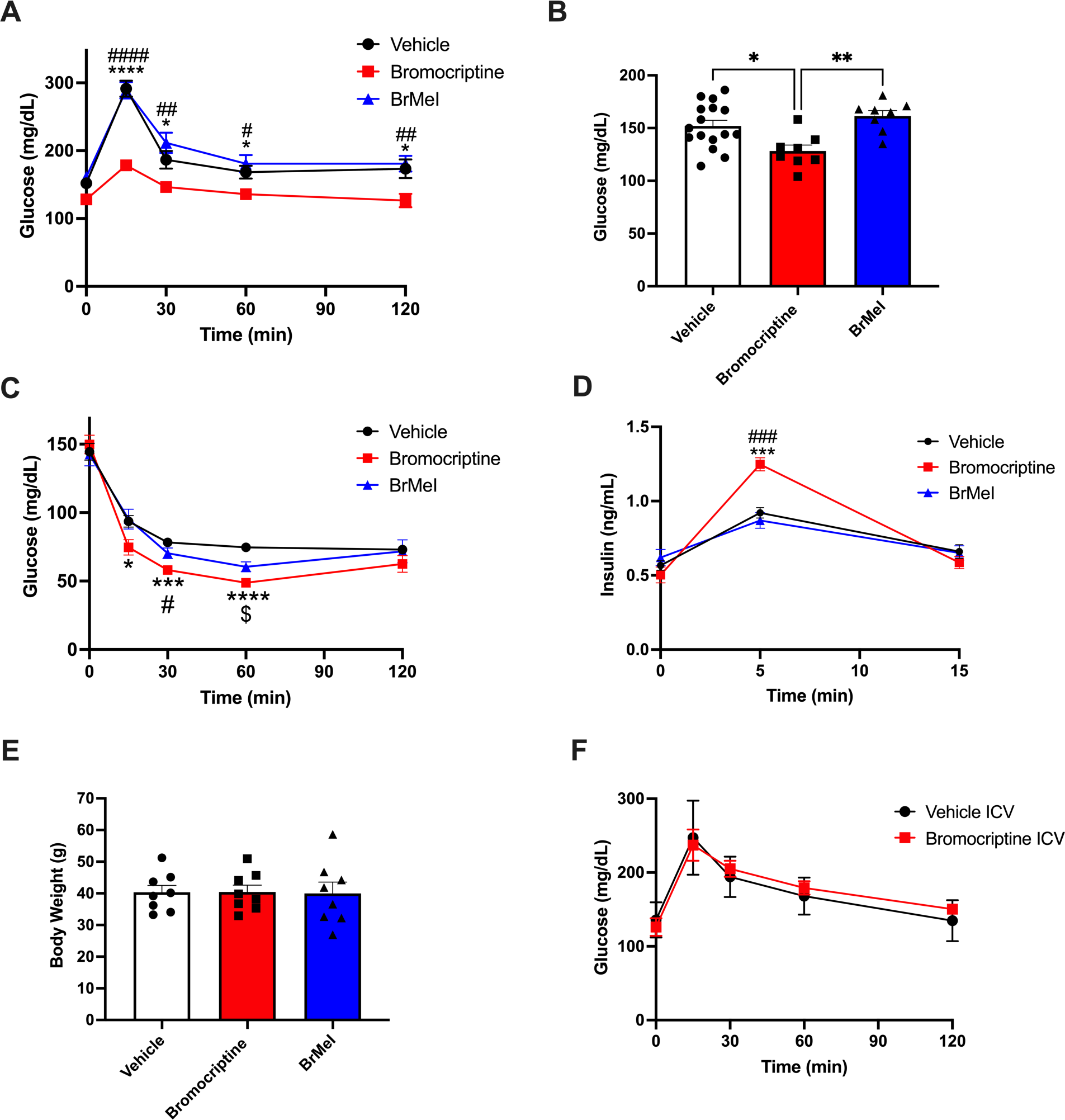
Tandem targeting of central and peripheral dopaminergic targets is required for treating dysglycemia *in vivo*. **(A)** Western diet-fed dysglycemic mice were systemically treated with bromocriptine (10 mg/kg i.p., in red), BrMeI (10 mg/kg i.p., in blue), or vehicle (i.p., in black). Oral glucose tolerance testing (OGTT, 2g/kg glucose) revealed significant effects of drug treatment (p < 0.0001) on improvement in glucose tolerance driven by bromocriptine (n = 8); BrMeI (n = 8) did not significantly modify glucose tolerance (p > 0.05) versus vehicle (n = 16). *<0.05 bromocriptine versus vehicle; #<0.05, ##<0.01, ####<0.0001 bromocriptine versus BrMeI. **(B)** Bromocriptine-treated mice (n = 8) exhibited significantly lower fasting blood glucose compared to BrMeI (p = 0.0035, n = 8) or vehicle (p = 0.0163, n = 16). *<0.05, **<0.01. **(C)** Insulin tolerance testing (ITT) showed that bromocriptine treatment (10 mg/kg i.p., in red; n = 8) led to significant improvements in insulin sensitivity versus vehicle (i.p., n = 8), with smaller improvements produced by BrMeI (10 mg/kg i.p., n = 8). *<0.05, ***<0.001, ****<0.0001 bromocriptine versus vehicle; #<0.05 bromocriptine versus BrMeI; $<0.05 BrMeI versus vehicle. **(D)** Bromocriptine treatment also significantly enhanced plasma insulin levels during the ITT compared to BrMeI (p = 0.0003) or vehicle (p = 0.0002) groups. ***<0.001 bromocriptine versus vehicle, ###<0.001 bromocriptine versus BrMeI. **(E)** There was no significant effect of bromocriptine or BrMeI on body weight versus vehicle treatment in Western diet-fed wild-type mice (p > 0.05) (n=8/group). **(F)** I.c.v. administration of bromocriptine (10 mg/kg, in red) in Western diet-fed dysglycemic wild-type mice. OGTT (2 g/kg glucose) did not significantly modify glucose tolerance versus i.c.v. vehicle treatment (in black) (n=8/group; p>0.05). All data represented as mean ± SEM.

Our *in vivo* metabolic results also raise the important question of whether the mechanisms responsible for systemic bromocriptine’s improvements in dysglycemia are through its actions on targets in the CNS versus those in the periphery or via its concurrent actions at both sites. To test whether bromocriptine’s effects are primarily due to CNS actions, we limited its actions to the brain via intracerebroventricular administration (i.c.v.) of the drug in Western-diet fed wild-type C57BL/6J mice. In contrast to systemic administration, i.c.v. bromocriptine had no significant effect on glucose tolerance (p > 0.05, **Figure 6F**). We validated these results by directly infusing DA precursor L-DOPA into the brain via i.c.v. administration in Sprague-Dawley rats for an extended 3.5-h period to permit adequate time for conversion into DA. Similar to our i.c.v. bromocriptine data, L-DOPA infusion did not significantly impact plasma glucose or insulin levels (p > 0.05; **Supplementary Figure 6**). Overall, these data suggest that both actions on targets in the CNS and periphery are required to effectively modify glycemic control.

## Discussion

Diabetes represents one of the most widespread conditions today, affecting ∼10% of the adult population of the United States^96^. Dysglycemia is central to the pathology of diabetes, resulting in a significant proportion of the morbidity and mortality associated with the illness^97, 98^. Treatment with APDs, some of the most widely prescribed psychiatric medications today, is a key contributor to the development of dysglycemia and T2D^99, 100^. Notably, though most of the focus on APDs’ metabolic symptoms has been on second-generation APDs like olanzapine^101, 102^, even first-generation drugs (*e.g.*, chlorpromazine and haloperidol) disrupt glycemic control, providing clues concerning the biological mechanisms underlying dysglycemia^26, 103–106^. Although multiple cellular targets for APDs have been identified^26, 37, 107–109^, the single unifying property of all APDs is their blockade of D2R and D3R. Conversely, bromocriptine, a D2R/D3R agonist can reduce dysglycemia and is FDA-approved to treat T2D^42, 110^. Together, this suggests an important role for DA and D2R/D3R signaling in glycemic control and treatment of dysglycemia.

DA’s emerging roles in metabolic regulation have been most extensively examined in the CNS. Significantly, both D2R and D3R are expressed in the hypothalamus, a brain region heavily involved in the homeostatic regulation of appetite and metabolism^17, 111, 112^. Indeed, hypothalamic D2R/D3R signaling mediates regulation of appetite and feeding behaviors that can result in compulsive overeating^17, 18, 113, 114^. Consistent with this, D2R blockade in the lateral hypothalamus increases food intake^17, 112, 115^. Hypothalamic D2R is expressed within arcuate nucleus neurons that are sensitive to peripheral hormone modulators of appetite and feeding including leptin and ghrelin, further linking central DA signaling to metabolic regulation, food intake and body weight changes, as well as to APD-related metabolic side effects^8, 10, 17, 19, 111, 116^. Moreover, hypothalamic D2R tonically regulates the release of prolactin, a hormone strongly associated with metabolic regulation as well as dysglycemia^117, 118^. There is also growing evidence of ongoing interplay between the CNS and periphery via leptin and ghrelin and dopaminergic mesolimbic and nigrostriatal circuits to modulate feeding^119–121^. In the striatum, D2R and D3R also modulate food-related reward, motivation, and anticipatory behaviors and are implicated in binge or compulsive eating^23, 122–124^. On the other hand, some reports suggest that CNS dopaminergic signaling in relation to feeding and reward is not sufficient to fully explain the mechanisms by which DA and D_2_-like receptors modulate metabolism^26, 125^. For example, changes in glucose homeostasis in response to D2R/D3R blockade by APDs occur even in the absence of increased food intake or psychiatric disease^126, 127^, and as little as a single administration of the APD olanzapine is sufficient to alter glucose homeostasis in healthy human subjects^33^. Recent discoveries that the endocrine pancreas uses local DA biosynthesis and signaling to modulate hormone release raises the intriguing possibility that dopaminergic regulation of metabolism may also rely on targets in the periphery^28, 32^. However, it has been very challenging to unravel central versus peripheral contributions of D2R/D3R signaling in metabolic regulation due to the current paucity of readily accessible pharmacological tools specifically targeting these respective compartments. Therefore, our goal was to generate a peripherally-limited D2R/D3R agonist to more selectively examine the metabolic roles of D2R/D3R signaling within the periphery. Such a drug could also serve as a comparator to the metabolic effects of D2R/D3R signaling in the CNS.

We employed a MeI quaternization strategy to render D_2_-like receptor-targeted drugs less BBB permeable. Indeed, MeI modification of compounds has been successfully used in experimental models of pruritus^128^, drug intoxication^77, 79^, substance dependence^129^ and withdrawal^78, 130^. We identified BrMeI as our lead compound based on its preserved receptor binding affinities for D2R and D3R as well as its efficacy in several *in vitro* functional assays, albeit with reduced potency. Consistent with this, higher concentrations of BrMeI shared bromocriptine’s receptor binding not only for D2R and D3R, but to additional GPCRs relevant to metabolic regulation including α_2A_ adrenergic and serotonin (5-HT_1A_, 5-HT_2A_, 5-HT_2C_) receptors^12, 26, 91, 92^. These receptors are well-established modulators of metabolism^32, 131–133^ and we posit that such additional targets contribute to bromocriptine’s therapeutic efficacy in improving dysglycemia.

Our *in vivo* metabolic data validates bromocriptine’s ability to improve dysglycemia within the established DIO model of T2D. This is consistent with extensive preclinical and clinical human data demonstrating bromocriptine’s efficacy in treating T2D dysglycemia^4, 134–137^. Work from our group and others points to bromocriptine’s actions in the periphery as an important component of its clinical effectiveness^12, 92^. We recently demonstrated that this drug also acts directly on peripheral dopaminergic targets including α-cell and α-cell D2R and D3R to reduce glucagon and insulin secretion^12^. Indeed, bromocriptine’s actions in the periphery may improve dysglycemia by: 1) diminishing hyperglucagonemia which in turn lowers hyperglycemia; and 2) decreasing chronic hyperinsulinemia. This induces a state of β-cell rest that resensitizes insulin-sensitive tissues and reduces cytotoxic β-cell stress^12, 32^. Beyond the endocrine pancreas, bromocriptine also acts on additional peripheral targets in adipose tissue including D_2_-like receptors and α_2A_ adrenergic receptors to further improve metabolism by modifying adipokine expression as well as diminishing adipogenesis and lipogenesis^138–140^. Collectively, these findings provide a strong rationale for our use of BrMeI to not only dissect central versus peripheral mechanisms of D2R/D3R-mediated glycemic control, but to potentially serve as a therapeutic tool capable of fine-tuning peripheral metabolism without potential CNS side effects.

According to our *in vivo* pharmacokinetic assay, BrMeI was preferentially distributed to the periphery shortly following systemic i.v. administration. In contrast to our original hypothesis, we discovered that BrMeI was equally distributed between the brain and periphery 1-hr post-administration. This suggests that the MeI modification did not fully block BrMeI’s entrance into the CNS over time. Nevertheless, compared to bromocriptine, BrMeI was present in the periphery as well as brain at much higher concentrations long after unmodified bromocriptine was cleared. Thus, despite its lower potency, BrMeI has a longer opportunity to signal, including in the periphery. Furthermore, BrMeI’s reduced efficacy in recruiting β-arrestin2 to D2R also suggests diminished D2R internalization and desensitization, enabling extended receptor signaling.

Our *in vivo* metabolic studies comparing the impacts of systemic administration of bromocriptine and BrMeI versus i.c.v. administration of bromocriptine and L-DOPA strongly suggest that stimulation of CNS dopaminergic targets by either bromocriptine or L-DOPA is insufficient to modify glycemic control. Instead, these data point to the importance of tandem targeting of both central and peripheral targets for effective glycemic control as well as for improvements in dysglycemia. Indeed, there is evidence of coordination between the hypothalamus and the autonomic nervous system (ANS) to control energy metabolism^141, 142^. Catecholamine signaling between the CNS and liver may be critical for this coordinated metabolic signaling. Bromocriptine-induced attenuation of hypothalamic noradrenergic drive, along with a reduction in the levels of circulating sympathetic mediators, diminished gluconeogenesis and lipolysis, provides an additional mechanism for improving dysglycemia^143, 144^. Ventromedial hypothalamic catecholamine activity was also correlated with a decrease in the hepatic transcription factors that potentiate gluconeogenesis and fatty acid oxidation after treatment with bromocriptine^5^. Finally, hypothalamic-ANS coordination mediating fat browning in relation to DA receptor agonism^3, 145, 146^ may explain the lack of net effects on weight or body composition with improved glucose tolerance (*i.e*., same fat mass, albeit with a different metabolic phenotype).

Limitations of the study include the incomplete limitation of BrMeI to the periphery with BBB penetrance over time. As a result, we cannot completely rule out that the BrMeI in the brain may contribute to the metabolic effects described here following its systemic administration. While possible, this is unlikely given the absence of significant effects in our OGTT studies. Despite sharing similar efficacies with unmodified bromocriptine in our GSIS and mitogenesis assays, it is also possible that BrMeI’s diminished potency may be responsible for the lack of an effective metabolic response. Evidence of a partial *in vivo* response in our ITT assay suggests that BrMeI retains some potency. Nevertheless, future work is clearly required to create the next generation of peripherally-limited D2R/D3R agonists with tighter exclusion from the brain across time. Another limitation is the lack of functional characterization of BrMeI’s actions on non-dopaminergic targets including serotonergic and adrenergic receptors. Lastly, our *in vivo* metabolic assays did not examine the impact of circadian rhythms on central and peripheral glycemic control by sampling at multiple zeitgebers, despite growing evidence of a strong circadian component to both central and peripheral dopaminergic metabolic regulation^122, 147–150^ – a topic for further studies.

Besides their utility in defining CNS versus peripheral contributions of the DA system to metabolic regulation, we propose that future generations of peripherally-limited D2R/D3R agonists can serve as especially effective treatments for APD-induced dysglycemia. Because APD-induced D2R/D3R antagonism is not limited to the CNS, these medications can act on peripheral dopaminergic targets to cause and/or exacerbate dysglycemia^26, 32, 37, 151^. As a result, co-administration of a D2R/D3R agonist whose actions are limited to the periphery would outcompete peripheral APD actions. Such a strategy would offset or overcome APD-induced dysglycemia with minimal risk of CNS/psychiatric side effects and without interfering with APDs’ intended therapeutic actions in the CNS.

Together, our results suggest that coordinated signaling via CNS and peripheral D_2_-like receptors is required for bromocriptine’s metabolic effects and underscores the importance of both peripheral and CNS dopaminergic metabolic regulation. Moreover, the design of peripherally-limited dopaminergic agonists opens the door to new classes of drugs for better, more effective treatment of dysglycemia.

## Data availability

All data will be made available by the corresponding author upon request.

## Author contributions

Synthesis and characterization of test chemicals were performed by AB, ME, CAB, JD, and AHN. Mitogenesis assays were performed by AJE and AJ. Insulin secretion assays were performed by ZJF. NanoBRET assays were performed by DA. Metabolic stability and pharmacokinetic studies were performed by RR and BSS. Rodent metabolic studies were performed by SP, MKH, and GJS. AB, JOMR, GJS, AHN, and ZF wrote the manuscript. Manuscript review and editing were performed by AB, ME, ZJF, DA, RR, SP, JOMR, CAB, BSS, AJE, AJ, MKH, GJS, AHN, and ZF. Project administration and supervision were provided by ZF.

## Acknowledgments

We gratefully thank Drs. Jeffrey Deschamps, Caitlin Burzynski, Jonathan Javitch, Vijay Yechoor, and George Gittes for assistance and helpful discussions throughout these studies. Funding support was provided by NIH grant R01DK124219 (ZF), K08DA031241 (ZF), DP1DA058385 (CAB), Department of Defense grants PR141292 (ZF, GJS), PR210207 (ZF), U.S. DOJ/DEA [Interagency agreement D-15-OD-0002] (AJ), Department of Veterans Affairs Merit Review [I01BX002758] and Department of Veterans Affairs Award Senior Research Career Scientist programs [1IK6BX005754] (AJ), and NIH/NIDA [Interagency agreement ADA12013] (AJ), the John F. and Nancy A. Emmerling Fund of The Pittsburgh Foundation (ZF), the NIDA Intramural Research Program Z1ADA000424 (AHN), the NIDA Medications Development Program (AHN), and the NIDA Addiction Treatment Discovery Program (ATDP). The contents do not represent the views of the U.S. Department of Veterans Affairs, U.S. Department of Justice, Drug Enforcement Administration, or the United States Government.

## Conflict of interest

The authors have declared that no conflict of interest exists.

## Notes

### Competing Interest Statement

The authors have declared no competing interest.

### Summary of Updates

This version of the manuscript has been updated to include an updated author list, updated author affiliations, updated manuscript, as well as an updated supplementary file.

