## Supplementary Information for "Development of novel tools for dissection of central versus peripheral dopamine D_2_-like receptor signaling in dysglycemia"

**A**

### Dopamine Receptor Binding

| BrMeI Concentration | Receptor / Radioligand | | | |
|---|---|---|---|---|
| | D1R % inhibition [$^3$H]SCH 23390 binding | D2R % inhibition [$^3$H]YM-09151-2 binding | D3R % inhibition [$^3$H]YM-09151-2 binding | D4R % inhibition [$^3$H]YM-09151-2 binding |
| 100 nM | 4.2 ± 7.6% | 13.3 ± 0.8% | 33.7 ± 3.7% | -2.2 ± 0.8% |
| 10 μM | 29.8 ± 6.1% | 92.2 ± 0.4% | 99.0 ± 0.1% | 47.1 ± 1.0% |

**B**

### Serotonin Receptor Binding

| BrMeI Concentration | Receptor / Radioligand | | |
|---|---|---|---|
| | 5-HT$_{1A}$ Receptor % inhibition [$^3$H]8OH-DPAT binding | 5-HT$_{2A}$ Receptor % inhibition [$^{125}$I]DOI binding | 5-HT$_{2C}$ Receptor % inhibition [$^{125}$I]DOI binding |
| 100 nM | 44.1 ± 4.0% | 21.7 ± 4.8%* | 4.7 ± 3.6%* |
| 10 μM | 99.4 ± 0.3% | 96.9 ± 0.6%* | 70.9 ± 1.7%* |

**C**

### Opioid Receptor Binding

| BrMeI Concentration | Receptor / Radioligand | | |
|---|---|---|---|
| | δ Opioid Receptor % inhibition [$^3$H]DPDPE binding | κ Opioid Receptor % inhibition [$^3$H]U69,593 binding | μ Opioid Receptor % inhibition [$^3$H]DAMGO Binding |
| 100 nM | 26.3 ± 3.8% | 15.6 ± 0.9% | -3.3 ± 1.5% |
| 10 μM | 59.5 ± 1.7% | 80.0 ± 11.0% | 72.0 ± 3.9% |

**D**

### Biogenic Amine Transporter Binding

| BrMeI Concentration | Transporter / Radioligand | | |
|---|---|---|---|
| | DAT % inhibition [$^{125}$I]RTI-55 binding | SERT % inhibition [$^{125}$I]RTI-55 binding | NET % inhibition [$^{125}$I]RTI-55 binding |
| 100 nM | 0.6 ± 8.8% | -6.5 ± 4.5%$^{\#}$ | 2.8 ± 8.3% |
| 10 μM | 35.4 ± 2.6% | 21 ± 11%$^{\#}$ | 37.0 ± 8.4% |

**Supplementary Figure 1. Bromocriptine methiodide receptor and biogenic amine transporter binding screen.** Bromocriptine methiodide (BrMeI) was screened via radioligand binding competition assays for binding to several families of G protein-coupled receptors as well as to biogenic amine transporters across two drug concentrations (100 nM, 10 μM). **(A)** Among DA receptors, 10 μM BrMeI showed significant binding to D2R and D3R, with 92.2% and 99.0% inhibition of specific radioligand binding, respectively; there was no significant D2R or D3R binding at 100 nM BrMeI. BrMeI showed no significant D1R or D4R binding at both assayed concentrations. **(B)** Among serotonin receptors, 10 μM BrMeI bound to 5-$HT_{1A}$, 5-$HT_{2A}$, and 5-$HT_{2C}$ receptors with 99.4%, 96.8%, and 70.9% inhibition of specific radioligand binding, respectively. There was no significant receptor binding inhibition with 100 nM BrMeI. **(C)** Among opioid receptors, 10 μM BrMeI showed binding to the δ-, κ-, and μ-opioid receptors with 59.5%, 80.0%, and 72.0% specific radioligand binding inhibition, respectively; there was no significant receptor binding by 100 nM BrMeI. **(D)** Among biogenic amine transporters, there was no significant BrMeI binding to the dopamine transporter (DAT), serotonin transporter (SERT) or norepinephrine transporter (NET) at either drug concentrations. Significant hits in the binding screen were defined as >50% inhibition of radioligand binding. Data are represented as % inhibition of specific control binding. Negative inhibition values indicate that more specific binding was measured in the presence of BrMeI than under control conditions. Unless otherwise noted, data represent the mean ± SEM performed in triplicate from n = 2 independent experiments. [*]Numbers represent the mean ± SEM performed in triplicate from n = 3 independent experiments. [#]Numbers represent the mean ± SEM performed in triplicate from n = 4 independent experiments.

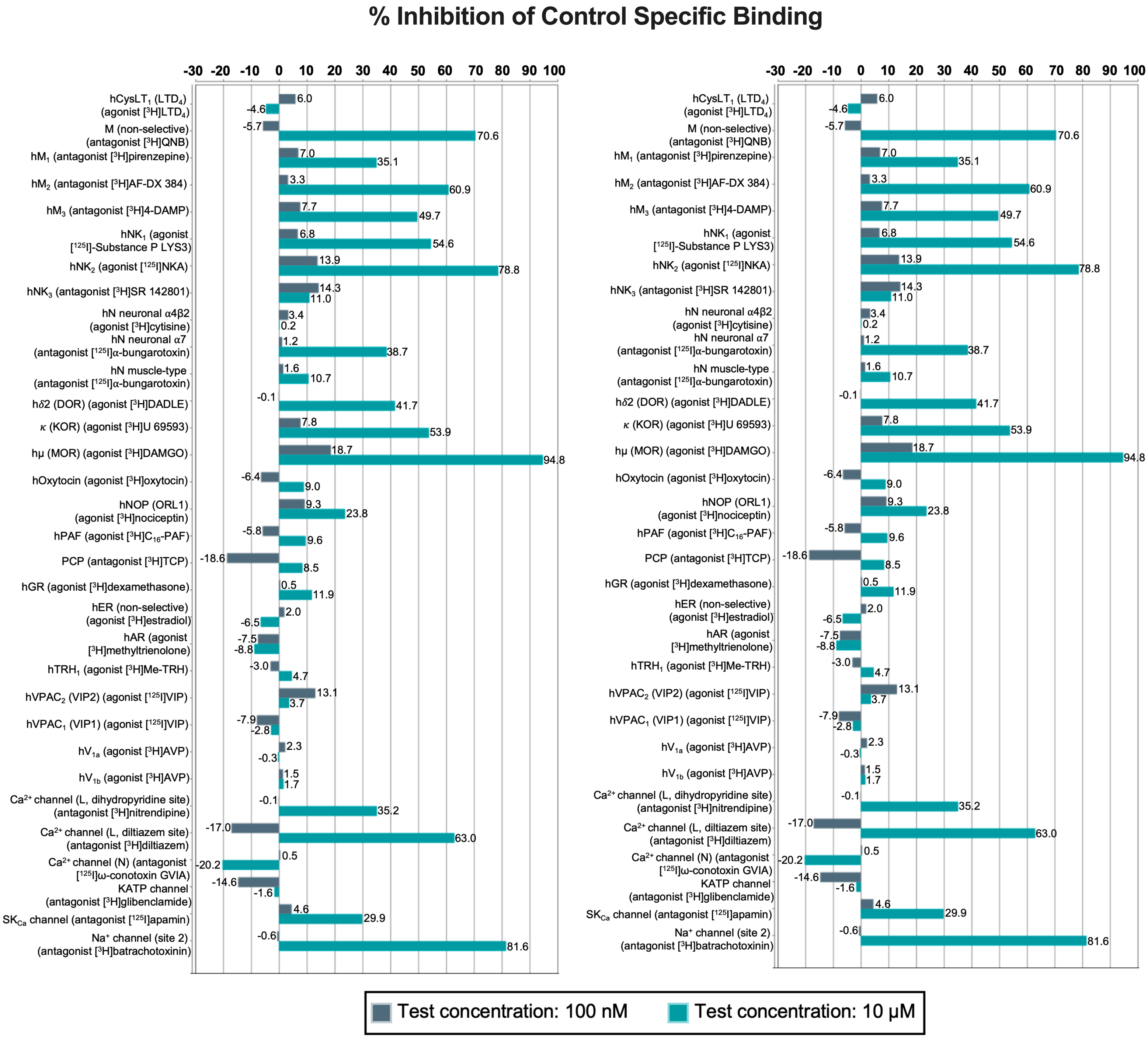


**Supplementary Figure 2. BrMeI off-target binding profile.** BrMeI was screened for off-target binding via radioligand binding competition assays across two drug concentrations: 100 nM (in grey), 10 μM (in blue). 100 nM BrMeI did not significantly bind screened targets. 10 μM BrMeI showed significant binding at $A_{2A}$, $\alpha_1$, $\alpha_2$, $D_{4.4}R$, κ-opioid, μ-opioid, M, $M_2$, $NK_1$, and $NK_2$ receptors as well as L-type $Ca^{2+}$ (diltiazem) and $Na^+$ (site 2) channels. Significant hits were defined as >50% inhibition of control specific radioligand binding; control radioligands are listed in parentheses. Data are represented as the mean % inhibition of specific control binding. Each experiment was conducted in duplicate.

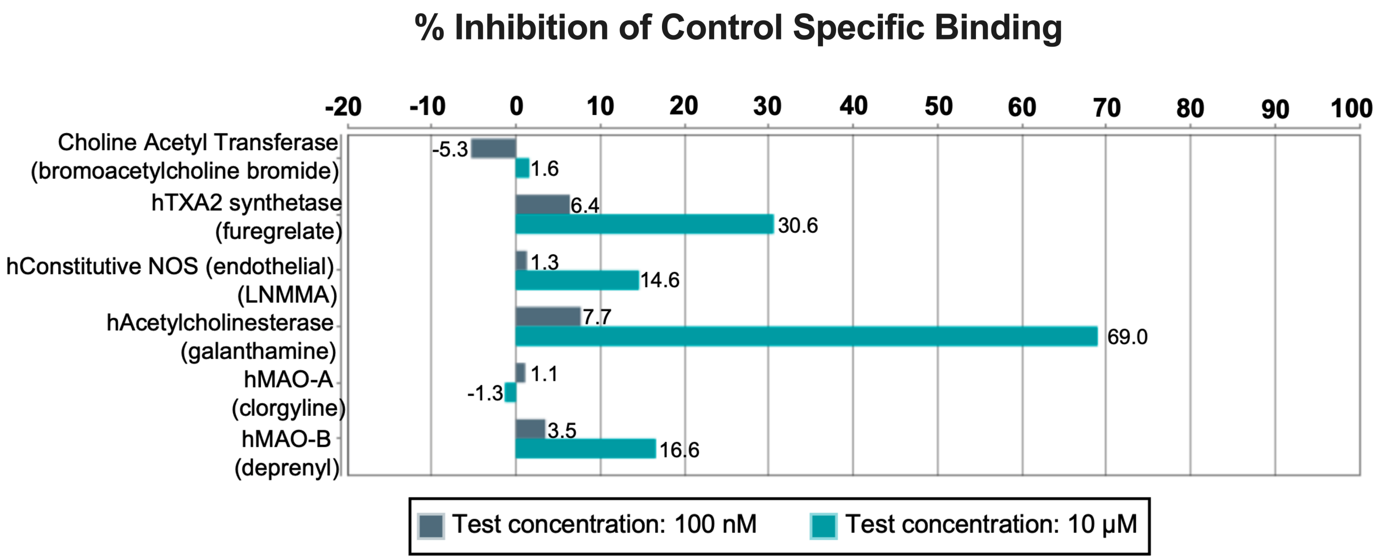


**Supplementary Figure 3. BrMeI off-target enzyme binding profile.** BrMeI was screened for off-target binding to several human enzymes using radioligand binding competition assays across two drug concentrations: 100 nM (in grey), 10 μM (in blue). 100 nM BrMeI demonstrated no significant binding. 10 μM BrMeI showed significant binding at acetylcholinesterase. Significant hits were defined as >50% inhibition of control specific radioligand binding; control radioligands are listed in parentheses. Data are represented as the mean % inhibition of specific control binding. Each experiment was conducted in duplicate.

**A**

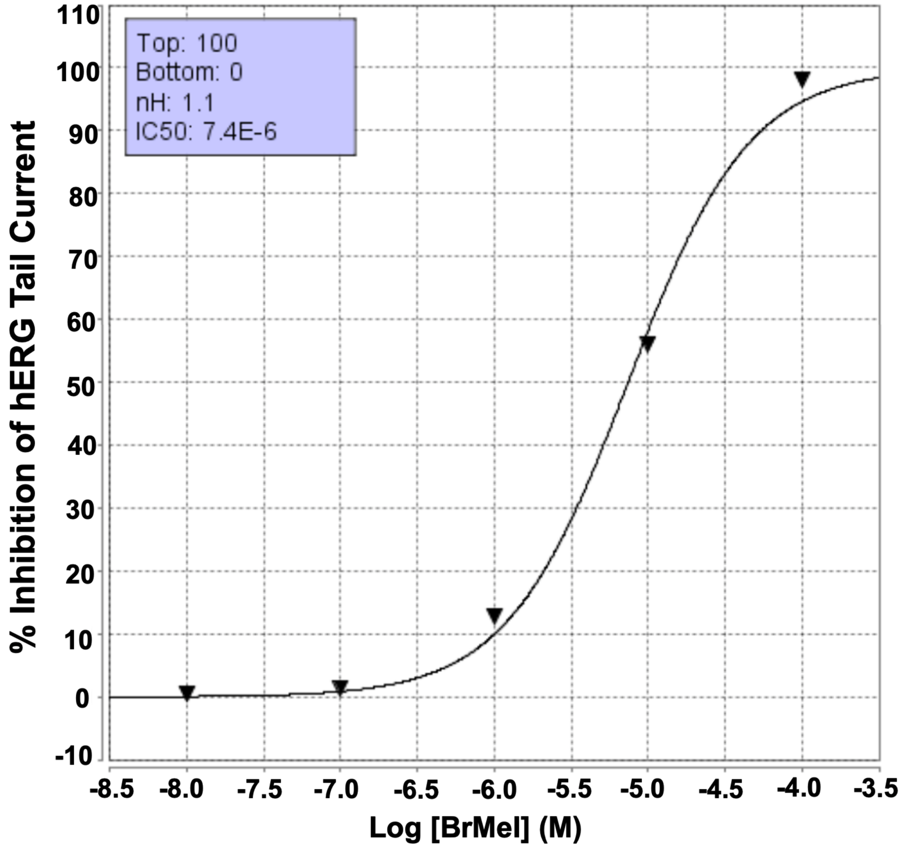


**B**

| hERG Assay | $IC_{50}$ (nM) | $IC_{50}$ Mean (nM) | $IC_{50}$ Range (nM) |
|---|---|---|---|
| Experiment 1 | 7400 | 8650 | 2500 |
| Experiment 2 | 9900 | | |

**Supplementary Figure 4. BrMel hERG channel assay.** BrMel was evaluated for hERG channel activity via measurement of hERG $K^+$ channel tail current amplitudes across a range of drug concentrations. **(A)** Representative BrMel dose response curve demonstrating concentration-dependent inhibition of hERG tail current ($IC_{50}$ = 7400 nM). **(B)** Quantification of $IC_{50}$ values from two separate hERG channel assays yielded a mean $IC_{50}$ of 8650 nM for BrMel. Data are represented as the % inhibition of hERG tail current, n = 2 independently conducted experiments.

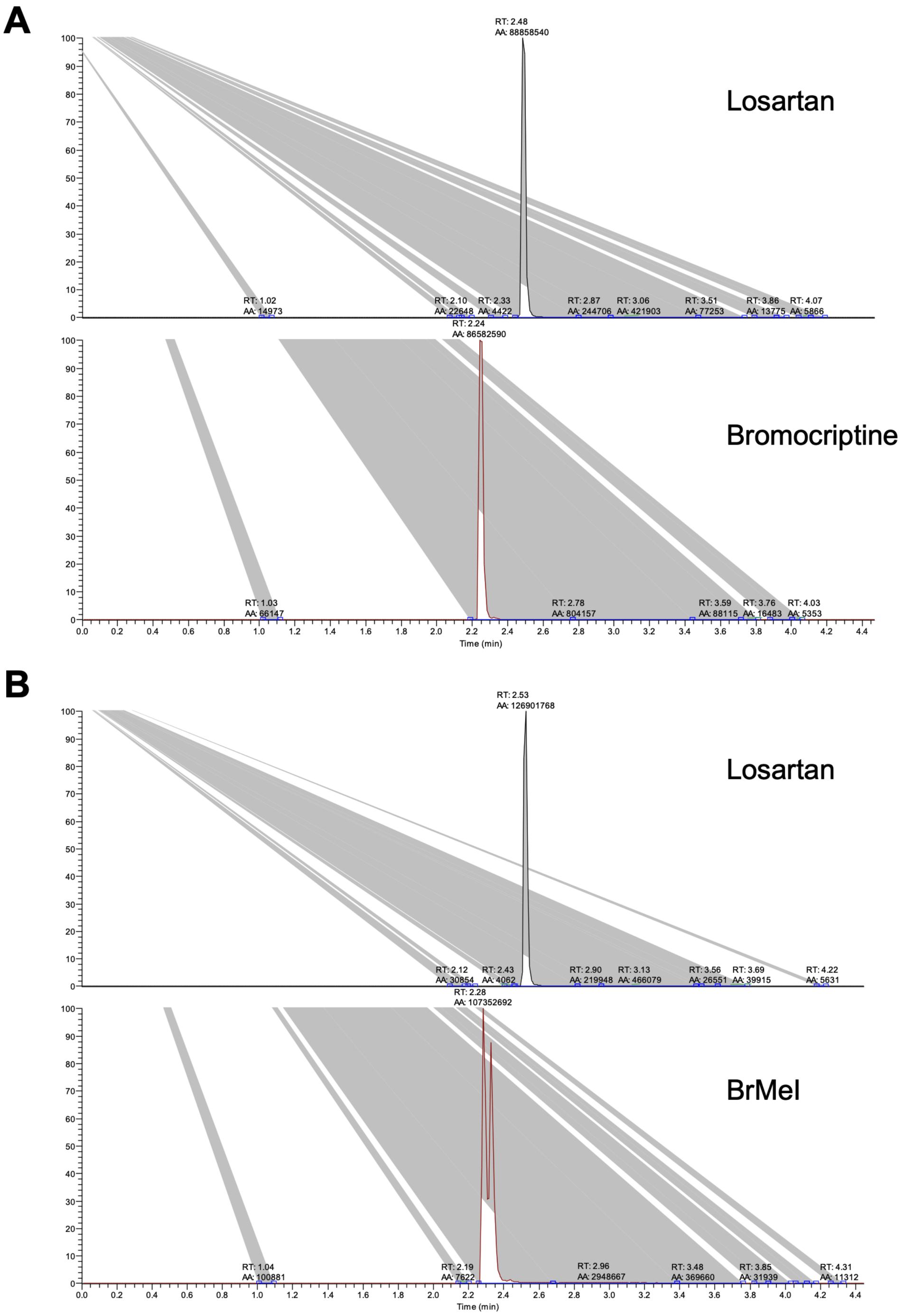


**Supplementary Figure 5. LC/MS/MS chromatography of BrMeI and bromocriptine in mouse pharmacokinetic assays. (A)** Representative chromatograms of internal standard losartan (upper panel) and bromocriptine (lower panel). **(B)** Representative chromatograms of losartan (upper panel) and BrMeI (lower panel).

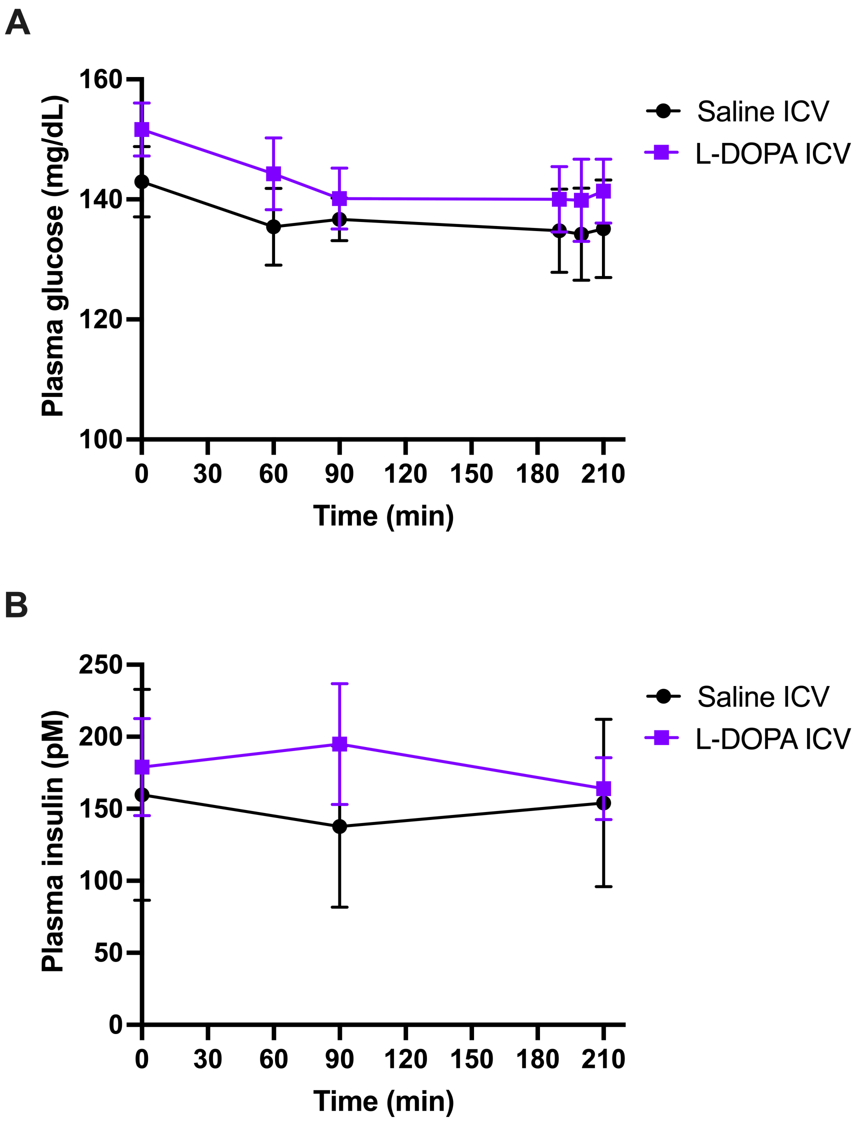


**Supplementary Figure 6. Stimulation of central dopaminergic targets is insufficient to modify plasma glucose and insulin levels in rats.** Direct infusion of DA precursor L-DOPA (10 mM, in purple; n = 4) i.c.v. for 3.5-h did not significantly modify plasma glucose **(A)** or plasma insulin **(B)** compared to the saline vehicle control (in black; n = 8) ($p > 0.05$). Data are represented as the mean ± SEM.

| Compound | Formula | Calculated | | | Found | | |
|---|---|---|---|---|---|---|---|
| | | C% | H% | N% | C% | H% | N% |
| **AB01-59** | $C_{14}H_{24}IN_3{\cdot}0.5H_2O$ | 45.41 | 6.81 | 11.35 | 45.56 | 6.50 | 11.11 |
| **AB01-60** | $C_{33}H_{43}BrIN_5O_5$ | 49.76 | 5.44 | 8.79 | 49.49 | 5.55 | 8.72 |
| **AB01-61** | $C_{24}H_{30}Cl_2IN_3O_2$ | 48.83 | 5.12 | 7.12 | 48.53 | 4.97 | 7.06 |
| **AB01-62** | $C_{17}H_{28}INO{\cdot}0.25H_2O$ | 51.85 | 7.29 | 3.56 | 51.92 | 7.14 | 3.44 |
| **MPE01-05** | $C_{13}H_{18}IN_3O{\cdot}H_2O$ | 41.39 | 5.34 | 11.14 | 41.08 | 4.91 | 11.11 |
| **MPE01-06** | $C_{21}H_{24}ClIN_2O$ | 52.24 | 5.01 | 5.80 | 51.95 | 4.96 | 5.66 |
| **AB01-117** | $C_{15}H_{22}INO_3{\cdot}1/3NH_4Cl{\cdot}1/3H_2O$ | 43.40 | 5.83 | 4.50 | 43.47 | 5.52 | 4.39 |
| **AB01-102** | $C_{21}H_{27}IN_2O$ | 56.01 | 6.04 | 6.02 | 55.66 | 6.12 | 6.02 |

**Supplementary Table 1. Elemental analysis of methiodide derivative compounds.**

| Compound | hD4R<br>$K_i$ ± SEM (nM) |
|---|---|
| **CAB03-015** | 1.50 ± 0.36 |
| **AB01-102** | 92.5 ± 21.1 |

**Supplementary Table 2. Radioligand competition D4R binding study data.** Binding affinities for human D4R as indicated by $K_i$ values were calculated for D4R-selective agonist CAB03-015 and its quaternary MeI analogue AB01-102. $K_i$ values are represented as the mean ± SEM with n≥3 independent experiments, each performed in triplicate.

| Compound | D4R | |
|---|---|---|
| | $EC_{50}$ ± SEM (nM) | $E_{max}$ |
| **CAB-03-015** | 0.24 ± 0.06 | 76.1% |
| **AB01-102** | 172 ± 59 | 71.4% |

**Supplementary Table 3. Functional activity in D4R-mediated adenylate cyclase/cAMP assay.** $EC_{50}$ and $E_{max}$ values were calculated for D4R-selective CAB-3-015 and its MeI analogue, AB01-102, from drug-stimulated dose-response curves in HEK cells expressing human D4.4 receptor (D4R) and adenylate cyclase type I. $EC_{50}$ values are represented as mean ± SEM. $E_{max}$ values represent % maximal efficacy relative to (-)-quinpirole stimulation. Data are from n≥3 independent experiments (CAB-03-015, n=6; AB01-102, n=10).

| Drug | $G\alpha_{i1}$ pEC$_{50}$ (EC$_{50}$, nM) | $G\alpha_{i1}$ $E_{max}$ | $G\alpha_{i1}$ pEC$_{50}$ (EC$_{50}$, nM) | β-arrestin2 $E_{max}$ |
|---|---|---|---|---|
| **DA** | 6.54 (288.67) | 100.00% | 4.56 (27634.23) | 100.00% |
| **Bromocriptine** | 7.82 (15.07) | 52.82% | 7.70 (20.12) | 28.00% |
| **BrMel** | 5.82 (1514.71) | 42.00% | 8.08 (8.37) | 24.32% |

**Supplementary Table 4. Potencies and efficacies of drug-stimulated G protein and β-arrestin2 recruitment to D2R.** Potency data are represented by pEC$_{50}$ with corresponding EC$_{50}$ values (in nM). $E_{max}$ values represent % maximal efficacy relative to DA treatment.

| Standards | Conc. tested | Dopamine Receptor/Radioligand | | | |
|---|---|---|---|---|---|
| | | **D1R % inhibition [$^3$H]SCH-23390 binding** | **D2R % inhibition [$^3$H]YM-09151-2 binding** | **D3R % inhibition [$^3$H]YM-09151-2 binding** | **D4.4R % inhibition [$^3$H]YM-09151-2 binding** |
| SCH-23390 | 100 nM | 98.88 ± 0.01% | | | |
| | 10 μM | 100.75 ± 0.15% | | | |
| Butaclamol | 100 nM | | 93.72 ± 0.03% | 93.30 ± 0.0% | |
| | 10 μM | | 102.32 ± 0.09% | 101.28 ± 0.06% | |
| Haloperidol | 100 nM | | | | 93.56 ± 0.31% |
| | 10 μM | | | | 100.65 ± 0.37% |

**Supplementary Table 5. Inhibition of specific DA receptor binding by standards.** Non-radiolabeled standards validated radioligand binding specificity at the respective DA receptors. Data are represented as the mean ± SEM performed in triplicate from n ≥ 2 independent experiments.

| Standards | Conc. tested | Serotonin Receptor/Radioligand | | |
|---|---|---|---|---|
| | | 5-HT1A % inhibition [$^3$H]8OH-DPAT binding | 5-HT2A % inhibition [$^{125}$I]DOI binding | 5-HT2C % inhibition [$^{125}$I]DOI binding |
| WAY 100,635 | 100 nM | 100.20 ± 0.26% | | |
| | 10 µM | 100.73 ± 0.22% | | |
| Ketanserin | 100 nM | | 78.92 ± 0.46% | |
| | 10 µM | | 96.28 ± 0.50% | |
| SB 242,084 | 100 nM | | | 98.80 ± 1.10% |
| | 10 µM | | | 99.56 ± 0.32% |

**Supplementary Table 6. Inhibition of specific serotonin receptor binding by standards.** Non-radiolabeled standards validated radioligand receptor binding specificity at the respective serotonin (5-HT) receptors. Data are represented as the mean ± SEM performed in triplicate from n ≥ 2 independent experiments.

| Standards | Conc. tested | Opioid Receptor/Radioligand | | |
|---|---|---|---|---|
| | | DOR % inhibition $[^3H]$DPDPE binding | KOR % inhibition $[^3H]$U69,593 binding | MOR % inhibition $[^3H]$DAMGO binding |
| Naltrexone | 100 nM | 65.70 ± 1.70% | | 85.60 ± 1.70% |
| | 10 µM | 95.37 ± 0.11% | | 99.67 ± 0.69% |
| Nor-BNI | 100 nM | | 97.67 ± 0.0% | |
| | 10 µM | | 100.17 ± 0.07% | |

**Supplementary Table 7. Inhibition of specific opioid receptor binding by standards.** Non-radiolabeled standards validated radioligand receptor binding specificity at the respective δ-opioid (DOR), $\kappa$-opioid (KOR), and µ-opioid (MOR) receptors. Data are represented as the mean ± SEM performed in triplicate from n ≥ 2 independent experiments.

| Standard | Concentration tested | Transporter / Radioligand | | |
|---|---|---|---|---|
| | | DAT % inhibition [$^{125}$I]RTI-55 binding | SERT % inhibition [$^{125}$I]RTI-55 binding | NET % inhibition [$^{125}$I]RTI-55 binding |
| Cocaine | 100 nM | 11.39 ± 0.60% | 6.1 ± 2.9% | 10.3 ± 2.7% |
| | 10 µM | 95.59 ± 0.59% | 94.2 ± 1.8% | 92.5 ± 1.7% |

**Supplementary Table 8. Inhibition of specific biogenic transporter binding by cocaine standard.** Non-radiolabeled cocaine validated biogenic amine transporter binding specificity at all three tested transporters. Data are represented as the mean ± SEM performed in triplicate from n = 2 independent experiments.
